# A fully defined pluripotent stem cell derived multi-liver-cell model for steatohepatitis and fibrosis

**DOI:** 10.1101/2020.09.03.280883

**Authors:** Manoj Kumar, Burak Toprakhisar, Matthias Van Haele, Asier Antoranz, Ruben Boon, Francois Chesnais, Jonathan De Smedt, Teresa Izuel Idoype, Marco Canella, Pierre Tilliole, Jolan De Boeck, Tine Tricot, Manmohan Bajaj, Adrian Ranga, Francesca Maria Bosisio, Tania Roskams, Leo A van Grunsven, Catherine M Verfaillie

## Abstract

Chronic liver injury, as observed in non-alcoholic steatohepatitis (NASH), progressive fibrosis, and cirrhosis, remains poorly treatable. Steatohepatitis causes hepatocyte loss in part by a direct lipotoxic insult, which is amplified by derangements in the non-parenchymal cellular (NPC) interactive network wherein hepatocytes reside, including, hepatic stellate cells, liver sinusoidal endothelial cells and liver macrophages. To create an in vitro culture model encompassing all these cells that allows studying liver steatosis, inflammation and fibrosis caused by NASH, we here developed a fully defined hydrogel microenvironment, termed hepatocyte maturation (HepMat) gel, that supports maturation and maintenance of pluripotent stem cell (PSC)-derived hepatocyte- and NPC-like cells for at least one month. The HepMat-based co-culture system modeled key molecular and functional features of TGFβ-induced liver fibrosis and fatty-acid induced inflammation and fibrosis better than monocultures its constituent cell populations. The novel co-culture system should open new avenues for studying mechanisms underlying liver steatosis, inflammation and fibrosis as well as for assessing drugs counteracting these effects.

## Introduction

Yearly approximately 2 million people die from liver disease worldwide, of which half are attributed to liver cirrhosis. Liver cirrhosis is an end-stage liver disease caused by long-standing injury resulting from among others, viral hepatitis, alcohol abuse, non-alcoholic steatohepatitis (NASH), or chronic exposure to chemicals (Asrani et al., 2019). Liver failure, occurring after long standing cirrhosis, is caused by loss of several critical functions of hepatocytes, including synthesis and secretion of plasma proteins, storage of biomolecules and micronutrients, regulation of glucose homeostasis, metabolism of drugs and blood detoxification. However, hepatocyte loss of function is caused not only by the direct viral or toxic insult to hepatocytes themselves, but is also significantly amplified by the disturbance of the cellular interactive network wherein hepatocytes reside (Marrone et al., 2016). This network involves the juxtaposition in the liver sinusoid of functionally interlinked non-parenchymal cell (NPC) types such as hepatic stellate cells (HSCs), liver sinusoidal endothelial cells (LSECs) and liver macrophages (Mϕs) (both liver resident Kupffer Cells (KCs), and peripheral blood Mϕs recruited to injured livers). Therefore, to study the mechanisms underlying liver insults, complex test systems incorporating multiple cell types would be required. Such models would also be needed to develop efficient therapeutic approaches that can reverse disease processes involving this interactive network.

To address how toxic insults affect hepatocytes as well as NPCs, murine or rat models are often used to study the effect of different hepatotoxins (Maes et al., 2016). However, interspecies differences between humans and rodents do not always allow extrapolating findings in animal models to the human patient. Therefore, co-cultures of human hepatoma cell lines with stellate cell, monocyte, and/or endothelial cell lines can be used (Asrani et al., 2019; Deng et al., 2019). In addition, and more relevant, co-cultures of primary human hepatocytes (PHHs) and NPCs can be stably maintained for several weeks (Baze et al., 2018; Messner et al., 2013). Although the latter best mimic human liver, the scarcity of human liver cells and high inter-donor variation (Hurrell et al., 2020; Kermanizadeh et al., 2019) represent important drawbacks.

With the advent of pluripotent stem cells (PSC) (Takahashi et al., 2007), it is now possible to create PSC-derived hepatocyte-(Corbett and Duncan, 2019), HSC-(Coll et al., 2018), EC-(Wilson et al., 2014) and Mϕ-like cells (Rajab et al., 2018). As PSCs are an inexhaustible cell population, this approach overcomes the issues related to low cell numbers and inter-donor variability of primary samples. Moreover, by using a collection of different iPSC lines, it would be possible to create models with known differences in susceptibility to liver damage. Some studies have co-cultured PSC-hepatocyte-like cell (HLC) progeny with mesenchymal cells and human umbilical vein endothelial cells (HUVECs) (Ayabe et al., 2018; Takebe et al., 2013). More recent studies created liver models by co-differentiation of PSCs to hepatocyte- and NPC-like cells directly in spheroids (Ouchi et al., 2019). An alternative is the assembly of pre-differentiated PSC-HLCs and -NPC-like cells into co-cultures (Koui et al., 2017). The latter approach might allow to recreate *in vitro* an model where HLCs and NPCs can be better numerically controlled compared to spontaneously co-differentiation of endodermal and mesodermal progeny.

We therefore here differentiated hepatoblast-, HSC-, endothelial-(ECs) and Mϕs-like cells from PSCs, that were then combined in a synthetic polyethylene glycol (PEG) hydrogel-based (Stevens et al., 2015) co-culture system. Using a design-of-experiment (DOE)-based screen we identified a unique combination of functionalizing extracellular matrix (ECM) component and cell-adhesion molecule (CAM) peptides (Lutolf and Hubbell, 2003), a metalloproteinase (MMP)-cleavable crosslinking peptide, and a hydrogel stiffness for optimal PSC-HLC maturation. We termed this hydrogel “hepatocyte maturation” or “HepMat” hydrogel. The HepMat hydrogel was shown to also support HSC-, EC- and Mϕ-like cells for at least one month. Finally, we demonstrated that the HepMat four-cell co-culture system was superior to HepMat monocultures for testing the ability of TGFβ or oleic acid (OA) to induce a fibrogenic and inflammatory cell phenotype, which could be blocked at least in part by treatment with obeticholic acid and to a lesser degree elafibranor.

## Results

### Combinatorial screen of instructive hydrogels to identify the optimal environment that sustains functional maturation of HLCs

We constructed a series of 3D hydrogels using four-arm-PEG building blocks with functional vinyl sulfone end-groups linked to different adhesion ligand peptides and matrix metalloproteinase (MMP) cleavable crosslinkers (Fig. 1 A). The MMP cleavable peptides, having thiol groups on both ends as described (Lutolf and Hubbell, 2003), were selected based on MMPs present in human livers (Duarte et al., 2015; Naim et al., 2017) and degradation kinetics named DG (degradability)1 (GCRD<u>VPLSYSG</u>DRCG), DG2 (GCRD<u>GPQGIAGQ</u>DRCG), and DG3 (GCRD<u>GPQGIWGQ</u>DRCG)(Patterson et al., 2010). To vary the hydrogel stiffness, three concentrations of PEG polymers were used named MP (mechanical property) 1 (8% w/v), MP2 (10% w/v) and MP3 (12% w/v). Bulk mechanical properties, measured using a nanoindenter, of the 8%, 10% and 12% w/v PEG hydrogels following peptide functionalization, were 3 Kpa, 9 Kpa and 20 Kpa (Fig. S1 A). For adhesion ligand functionalization, we selected 24 peptides representing the active part of ECM components (fibronectin, collagen I, collagen III, collagen IV, laminins, perlecan) or cell adhesion molecules (E- and N-cadherin) present in liver (based on data from the human Matrisome project (Naba et al., 2016)(Table 1). The peptide sequences consisted of Ac-G**C**GYGXXXXG-NH_2_ where ‘XXXX’ represented the active component of the peptide. The efficient binding of the ECM/CAM peptides to the PEG backbone was demonstrated by fluorescence intensity measurements of hydrogels conjugated with different concentrations of a fluorescently labeled peptide (Fig. S1 B).

**Table 1:**
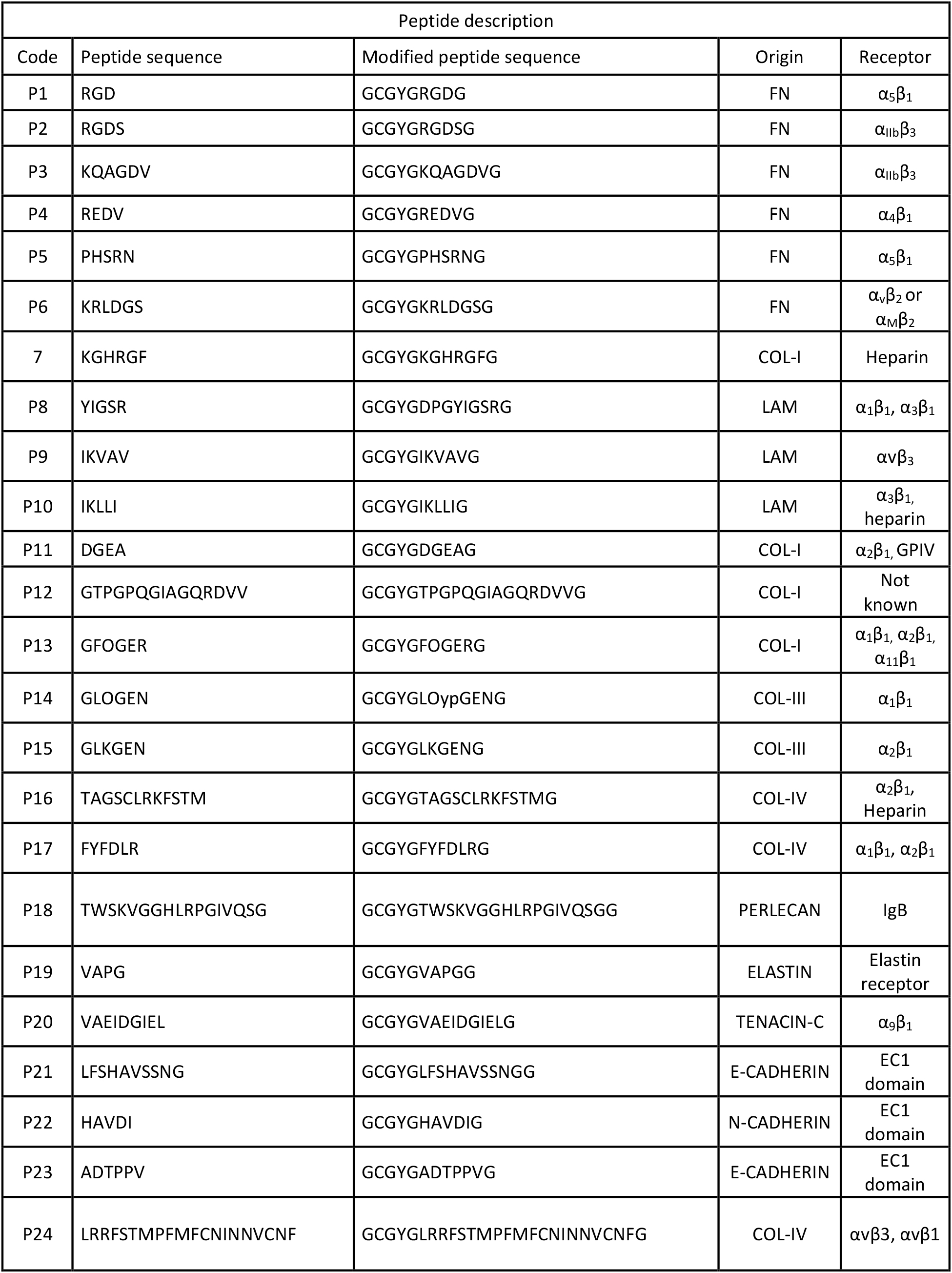
ECM peptide sequences

**Figure 1.**
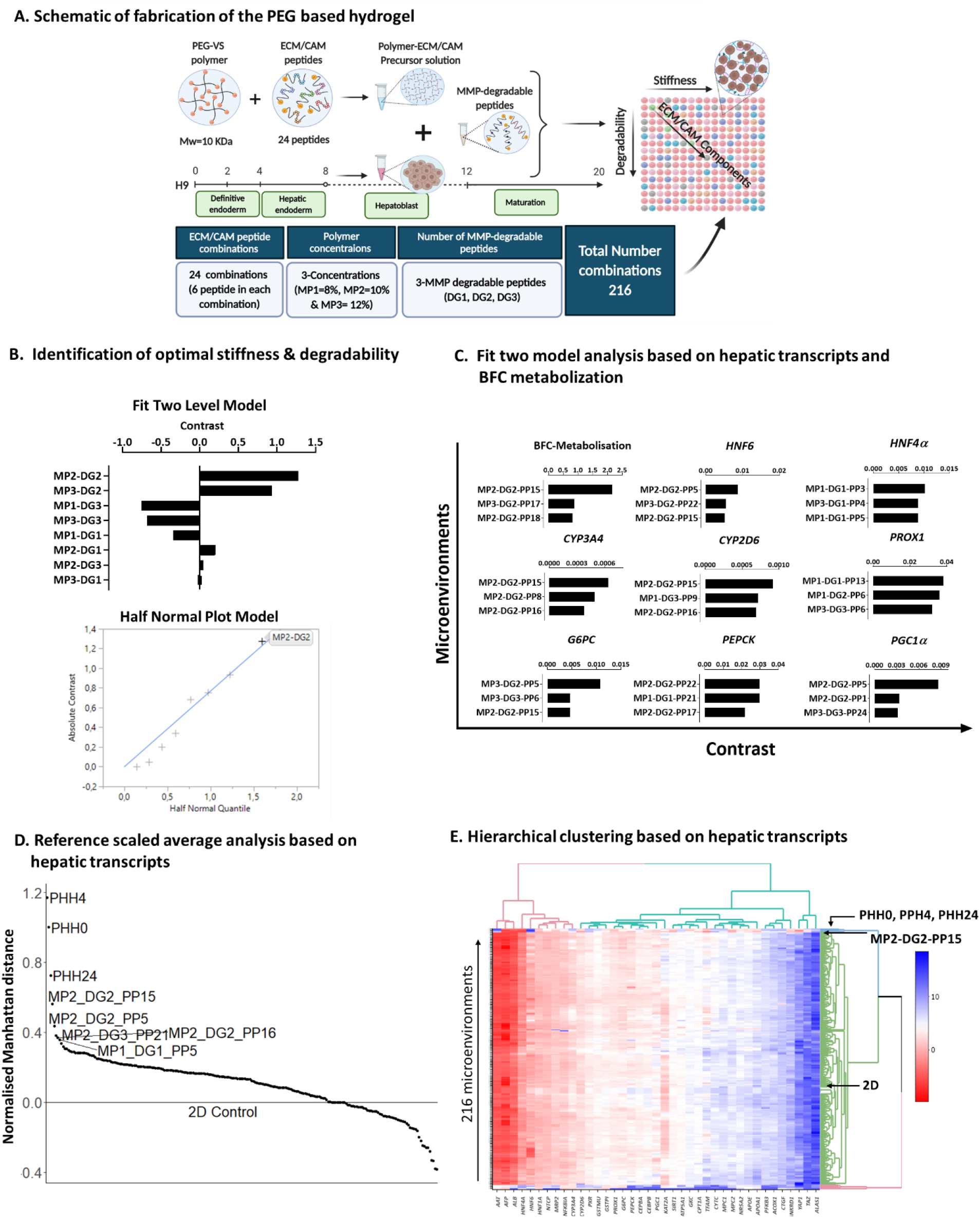
Identification of optimal mechanical properties and peptide functionalization of polyethylene glycol (PEG) hydrogels to support maturation of PSC-hepatocyte like cells (HLCs). **(A)** Schematic of PEG based hydrogel fabrication with PSC-liver hepatoblasts and 20-day PSC-HLC differentiation protocol. **(B)** Analysis of DOE experiment using the Fit Two Level Model- and Half Normal Plot Model-based analysis from the JMP-pro package based on transcript levels for 8 hepatocyte genes (*HNF4α, HNF6, PROX1, CYP3A4, CYP2D6, PEPCK, G6PC, PGC1α*) and BFC metabolization to define the degradability and stiffness of the hydrogel that most optimally supports HLC maturation. **(C-D)** Analysis of DOE to identify the hydrogel characteristics (peptide pool, MMP linker and PEG concentration) that supports the highest maturation of PSC-HLCs (C) Fit Two Level Model based on transcript levels for 8 hepatocyte genes and BFC metabolization. (D) Reference Scaled Average analysis based on 8 hepatocyte gene expression levels. **(E)** Hierarchical clustering analysis based on transcript levels of 39 hepatocyte marker genes (see table 3) of day 20 PSC-HLC progeny in the 216 hydrogels, compared with PHH0, PHH4 and PHH24 and HLCs in 2D culture. Cluster gram shows delta CT compared with the *RPL19* housekeeping transcripts; red is higher expression and blue lower expression. MP1, MP2, MP3 = mechanical property based on different PEG concentrations; DG1, DG2, DG3 = degradation property, based on different MMP cleavable linkers; DOE = design of experiment; BFC = 7-benzyloxy-4-trifluoromethylcoumarin; PHH = primary human hepatocytes

To define the environment that most optimally sustained PSC-HLCs and to avoid prohibitively exhaustive screening, we used a discrete numeric design (using JMP pro (SAS)) to create a design-of-experiment (DOE) (Li et al., 2012). This resulted in a total of 216 microenvironments, each consisting of one of three stiffnesses (MP1-3), one of three MMP linkers (DG1-3), and pools of 6 peptides (PP1-PP24) (Table 2). To test the effect of these microenvironments on HLC maturation, we seeded PSC-progeny differentiated to the hepatic lineage for 8 days in 2D culture (hepatoblast stage) at 3×10^5^ cells/10 μL of the different hydrogel compositions, and allowed the cells to mature for an additional 12 days. Maturation was assessed by RT-qPCR for mature hepatocyte marker genes (Table 3) and benzyloxy-4-trifluoromethylcoumarin (BFC) metabolization (function of CYP3A4 (and CYP1A2)). All data was compared with HLCs maintained until day 20 in 2D culture, and PHHs (freshly thawed [PHH0]; cultured for 4h [PHH4], and 24h [PHH24]).

**Table 2:**
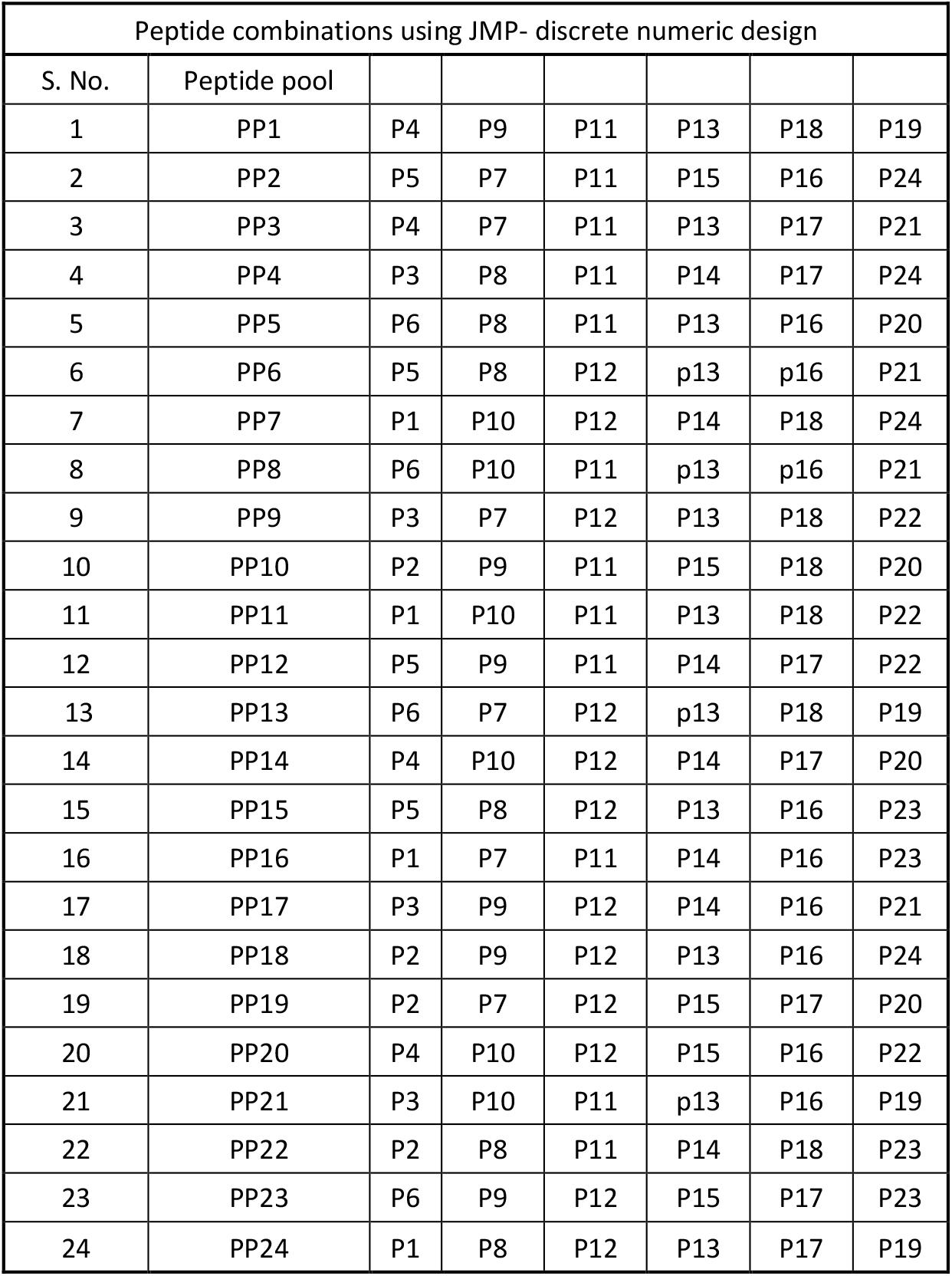
Peptide pools included in the DOE

**Table 3.**
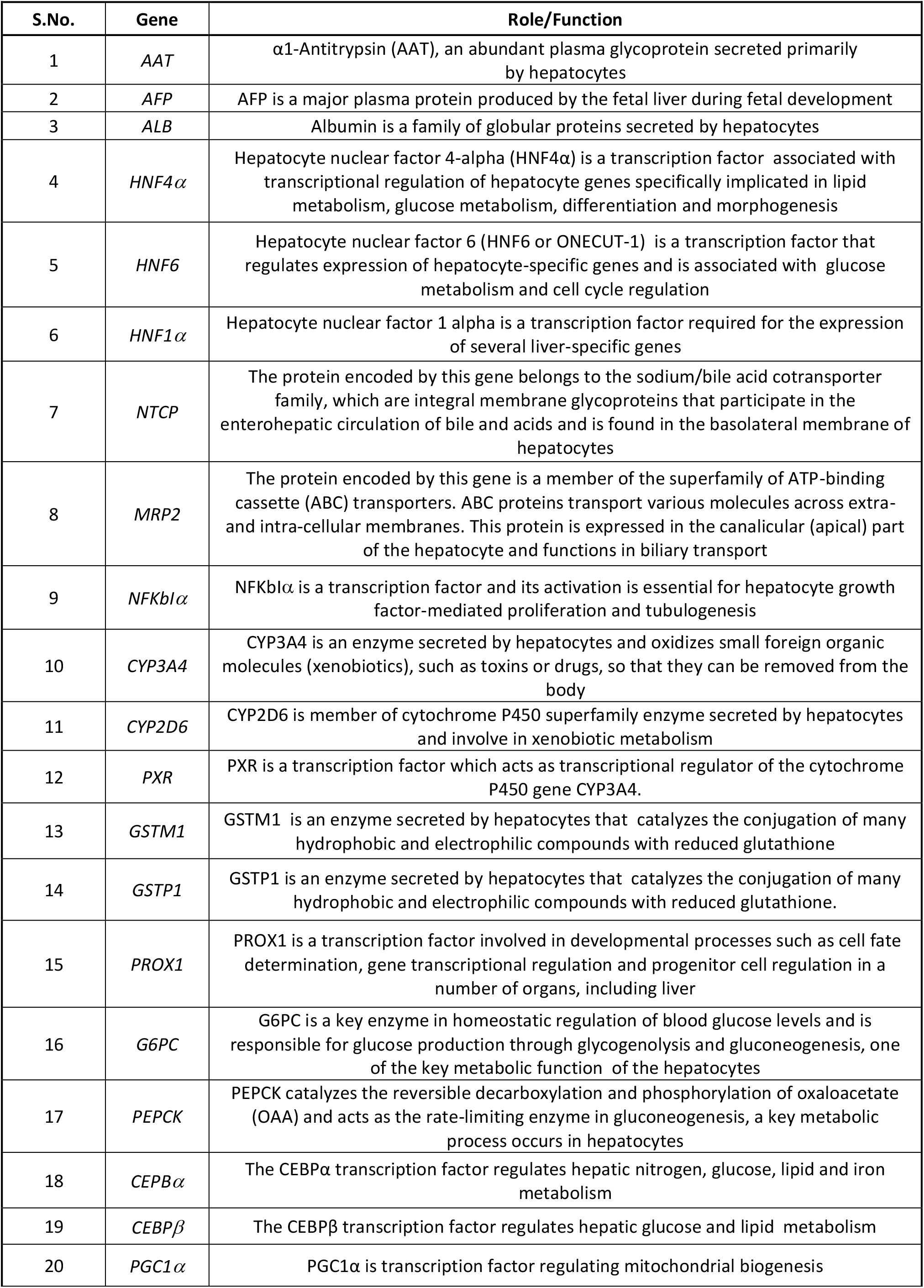
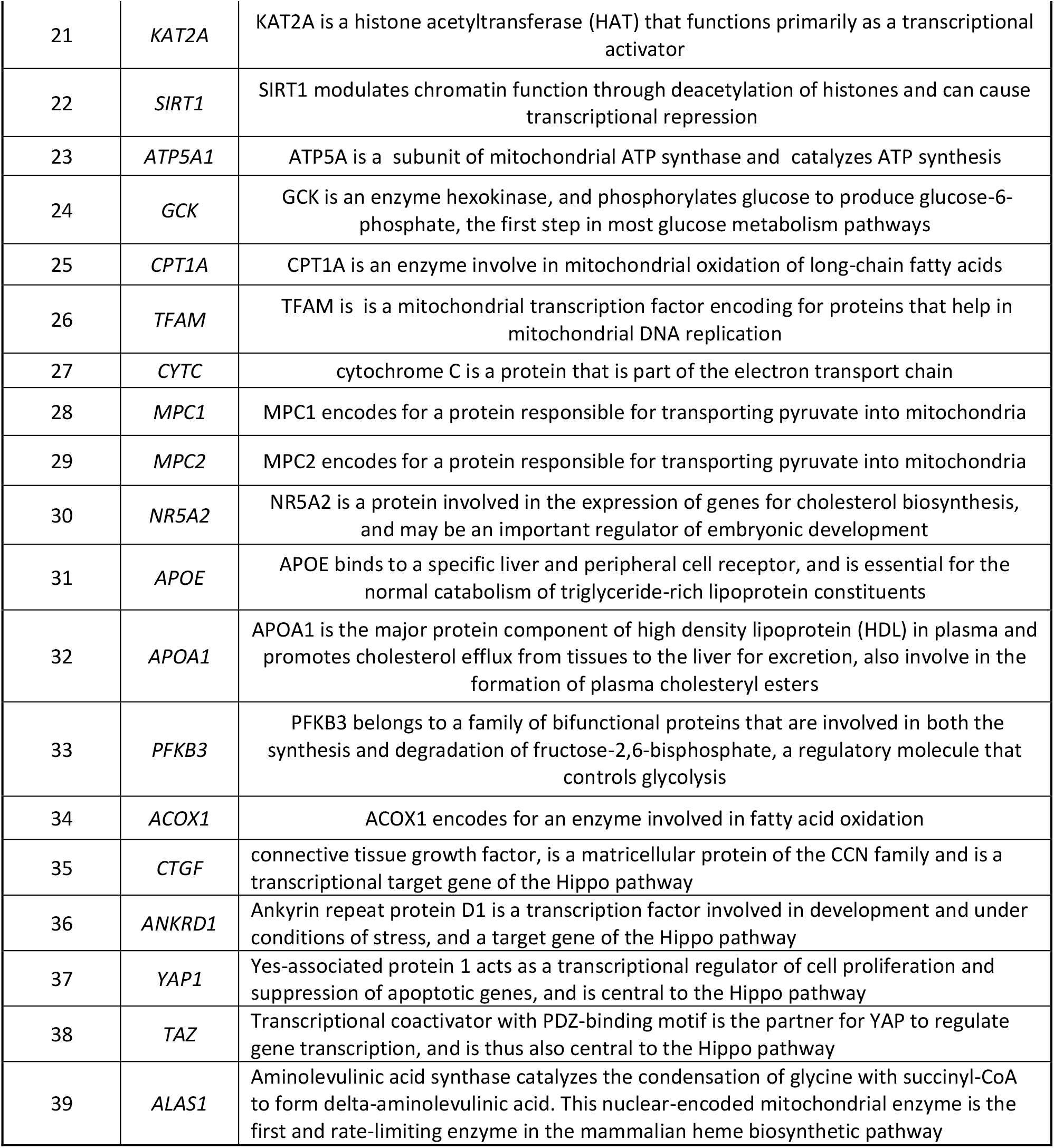
Hepatic gene transcripts tested

Using the Fit-Two-Level model screening module, which identifies effects that have a large impact on the response based on the sparsity-of-effects principle (Box and Meyer, 1986), we identified that among gels with different mechanical properties (i.e. the combinatorial effect of DG1, DG2 or DG3 with MP1, MP2 or MP3) the combination of MP2 (9 Kpa) and DG2 (GCRD<u>GPQGIAGQ</u>DRCG) yielded hepatic progeny with the greatest maturity (RT-qPCR for 8 hepatocyte genes and BFC metabolization). This was confirmed using a half-normal probability plot, which estimates the effect of a given main effect or interaction and its rank relative to other main effects and interactions, enabling the ranking of factors by importance (Fig. 1 B). We next analyzed the combined effect of mechanical properties and functionalization by different peptide pools on hepatocyte maturation, using the same 8 hepatic transcript levels and BFC metabolization as readout. The Fit-Two-Model module identified that MP2-DG2 hydrogels functionalized with peptide pool 15 (PP15) ranked among the top 3 microenvironments for five out of nine 5/9 markers tested (Fig. 1 C). Calculation of normalized Manhattan distances (based on log2 fold hepatic gene expression changes) revealed that hepatic progeny from the MP2-DG2-PP15 hydrogel-based cultures ranked the closest to PHHs (Fig. 1 D). Finally, hierarchical clustering based on 39 different mature hepatocyte gene marker transcripts (Table 3) also demonstrated that the hepatic progeny from MP2-DG2-PP15 hydrogel-based cultures ranked closest to PHHs (Fig. 1 E).

To confirm the optimal peptide pool for HLC maturation, we repeated the screen (Fig. S1 C-E) starting from the MP2-DG2 backbone and functionalized with any of the 24 different peptide pools. Hierarchical clustering analysis of RT-qPCR results from day 20 progeny identified again the PP15 peptide pool functionalization of the MP2-DG2 hydrogel to be the most supportive for HLC maturation. Therefore, we concluded that the peptide combination PP15, containing the fibronectin peptide P5, the laminin peptide P8, the collagen I peptides P12 and P13, the collagen IV peptide P16 and the E-cadherin peptide P23 (Table S1), in 10% PEG-hydrogels crosslinked with the GCRD<u>GPQGIAGQ</u>DRCG-MMP9 degradable linker, generates PSC-HLCs that have transcriptional and functional characteristics that align most closely with PHHs.

### The hepatocyte maturation (HepMat) hydrogel also supports further maturation of HLCs generated from genome engineered PSC (HC3x-PSC) and cultured in amino-acid rich (AAGly) medium

In a recent study (Boon et al., 2020), we demonstrated that PSC-hepatic differentiation is significantly enhanced when PSCs are genetically engineered to inducibly overexpress three transcription factors (*HNF1A, FOXA3* and *PROX1*, termed HC3x-PSC) from day 4 onwards, and are cultured in amino acid (AA)-enriched medium (liver differentiation medium [LDM] supplemented with 16 ml non-essential AA solution, 8ml of essential AA solution per 100 ml of LDM, and 20 g/l glycine; termed AAGly medium).

We first determined if the MP2-DG2-PP15 hydrogel is also the most optimal hydrogel composition to induce maturation of HLCs when cultured in AAGly medium. As shown in Fig. S1 D&E, among the different peptide pool combinations, the MP2-DG2-PP15 (HepMat) hydrogel-cultured HLCs again aligned most closely to PHH. We next embedded HC3x genome engineered hepatoblasts in HepMat hydrogels and cultured them for 40 days in AAGly medium, conditions shown previously to induce the greatest HLC maturation (Boon et al., 2020). RT-qPCR analysis demonstrated that HC3x progeny, 32 days after embedding in HepMat hydrogels and maintained in AAGly medium, expressed significantly higher transcript levels for all marker genes tested compared with HC3x-HLCs cultured for 40 days in 2D in AAGly medium (Fig. 2 B). Moreover, CYP3A4 activity of HC3x-HLCs, already high in AAGly cultured 2D HC3x progeny (as also described by (Boon et al., 2020)), increased a further 4±0.3-fold when HC3x-HLCs were cultured in 3D-HepMat-AAGly conditions (Fig. 2 C). When 3D-HepMat-AAGly HC3x-HLCs were treated with rifampicin, BFC metabolization increased by 9.9±1.3 fold. No significant differences were observed in albumin secretion between 2D-AAGly and 3D-HepMat-AAGly cultured HC3x-HLCs (Fig. 2 C). Interestingly, although a significant increase in transcript levels for *G6PC and PGC1α* was also seen when HC3x cells were cultured in 3D-HepMat-LDM conditions between day 12 and 40 compared with 40 days in 2D LDM-culture, the cytochrome transcript levels were not induced (Fig. S2 A). Thus, in line with what we published for 2D cultures, AA levels in the culture medium are important for hepatocyte maturation also in 3D-HepMat cultures.

**Figure 2.**
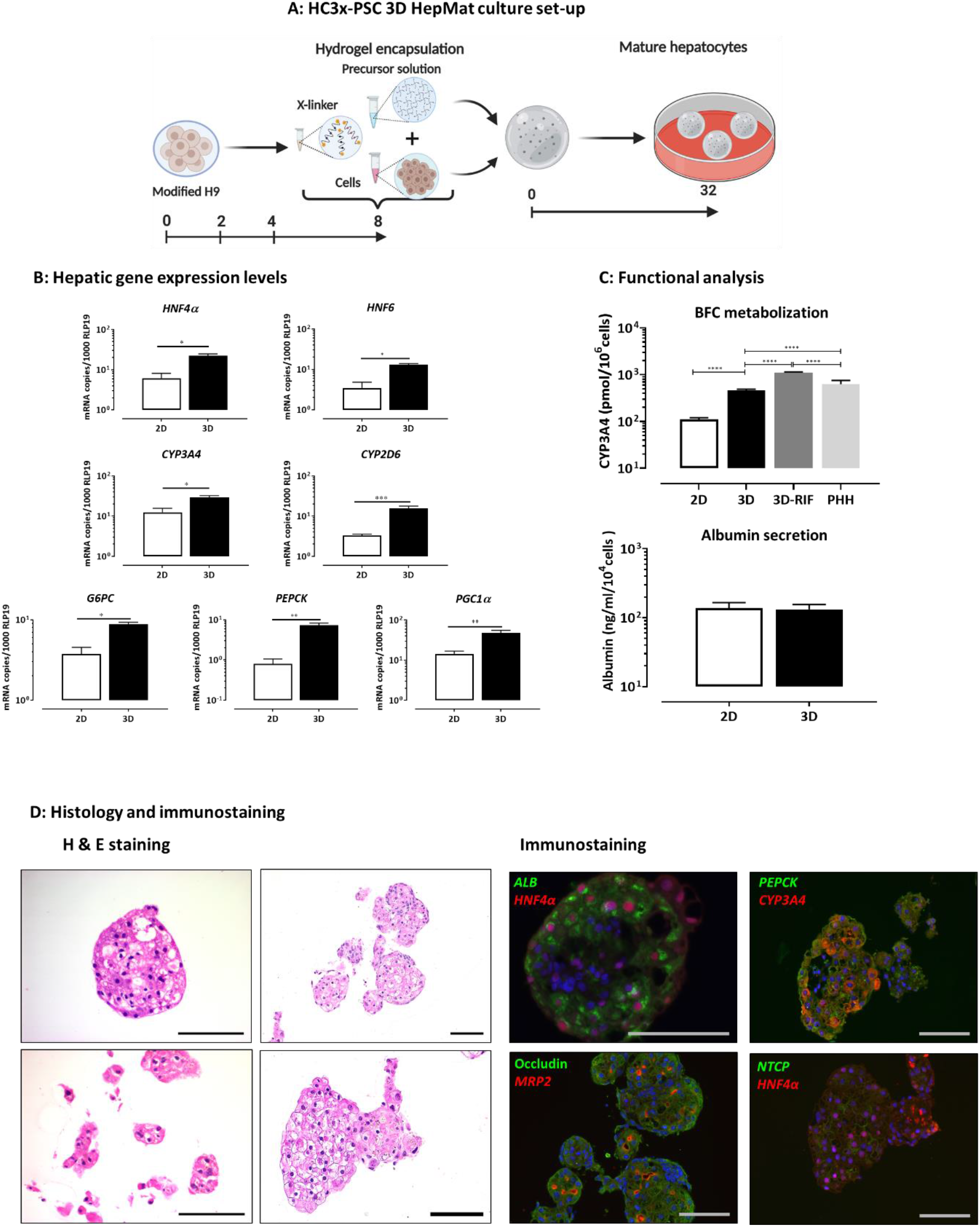
The HepMat hydrogel supports further maturation of PSC-HLC. **(A)** Schematic representation of culture of genome engineered HC3x-PSCs initially for 8 days in 2D culture and then in HepMat hydrogels for 32 days using liver differentiation medium + extra amino acids (AAGly) from day 14 onwards as described (Boon et al., 2020). (**B)** Gene expression of *HNF4α, HNF6, CYP3A4, CYP2D6, G6PC, PEPCK and PGC1α* in HC3x-progeny maintained in 2D culture for 40 days vs. progeny of d8 2D-culture derived hepatoblasts embedded in HepMat hydrogels for an additional 32 days (N=3 biological replicates). (**C)** Functional analysis of day 40 HC3x-progeny in 2D or 3D-HepMat cultures: Upper panel: CYP3A4 function defined by BFC metabolization (2D: N=5; 3D: N=13; 3D-RIF: same as 3D but cells were exposed to 25μM of Rifampicin (RIF) for 48h, N=4; PHH, N=3; N = biological replicates); Lower panel: Albumin secretion (N=3 biological replicates). (**D)** Histology and immunostaining of day 40 HC3x-progeny in 3D HepMat cultures: Left panel: H & E staining of cells; Right panel: Immunostaining for albumin & HNF4α, CYP3A4 & PEPCK, MRP2 & Occludin and NTCP & HNF4α (Representative example of 3 biological replicates). Data are shown as mean ± SEM and analyzed by two –tailed student t-test (RT-qPCR) or one way Anova (Tukey’s multiple comparison for BFC). *p <0.05; **p <0.01; Scale bars= 100 μm; PHH = primary human hepatocytes.

Histological analysis of the HC3x progeny cultured in 3D HepMat-AAGly identified polygonal cells with an eosinophilic and clear/vacuolated cytoplasm, with a round shaped nucleus, and present in a cohesive pattern, highly reminiscent of hepatocytes. (Fig. 2 D). Immunostaining and confocal imaging confirmed the hepatocyte identity of the cells (nearly 100% albumin and HNF4α positive) (Fig. 2 D). We could also identify CYP3A4 and PEPCK positive cells. Interestingly, there was little overlap between CYP3A4 and PEPCK staining (Fig. 2 D), consistent with hepatocyte diversity in primary human liver (MacParland et al., 2018). In addition, most cells had cell surface staining of NTCP, and stained for MRP2 both in the cell cytoplasm and more punctate on the cell membrane.

### The hepatocyte maturation hydrogel also maintains of PSC-derived non-parenchymal cells

As hepatocytes are located in close proximity with NPCs in liver sinusoids *in vivo*, we hypothesized that the HepMat hydrogel that supports HLCs should also maintain PSC-derived NPCs. We therefore embedded PSC-EC, PSC-HSC and PSC-Mϕ progeny harvested from 2D cultures in HepMat hydrogel in their respective culture media, and maintained the 3D cultures for 32 days. Survival and phenotype were assessed by RT-qPCR, immunostaining or flow cytometry, and functional studies.

To generate ECs, we used PSCs wherein the master-regulator, *ETV2*, was incorporated in the safe harbor locus, *AAVS1*, under the control of a TET-ON promoter (iETV2-PSC) (Elcheva et al., 2014; Ordovas et al., 2015). The *ETV2* transcription factor was induced with doxycycline from day 0 of differentiation, and doxycycline maintained for the duration of the culture. By flow cytometry, iETV2-PSC progeny on day 8 was nearly 100% CD31 and KDR double positive (Fig. S3 A). RT-qPCR demonstrated that 2D-cultured day 8 and 12 iETV2 progeny expressed high levels of *CD31*, but very low levels of LSEC-specific genes (*FGFR2B, STAB1*, and *CLEC4G;* and the recently defined LSEC markers *FCN3, OIT3, CLEC4M* (Ramachandran et al., 2019) (Fig. 3 A and S3 A). iETV2-ECs harvested on day 12 from 2D cultures were then embedded in HepMat hydrogel, and maintained in EC medium with doxycycline, bFGF and fetal bovine serum (FBS). iETV2-ECs remained viable until day 32 (calcein-AM staining). iETV2-ECs formed tubes 5-6 days after embedding in the gels. However, these tubes disintegrated within 2-3 days (Fig. 3 A, left panel of bright field imaging). When the medium was supplemented with endothelial cell growth supplement (ECGS), tube formation in HepMat hydrogels was stable for ±2 weeks (Fig. 3 A right panel of bright field imaging). Nearly 100% of d32 HepMat-cultured iETV2-ECs were VE-Cadherin positive (Fig. 3 A). RT-qPCR analysis 32 days after embedding iETV2-ECs in HepMat hydrogels demonstrated persistent expression of *CD31*, and a modest induction of some (*FCGR2B, STAB1, LYVE1* and *FCN3*) but not all (*OIT3, MRC1, CLEC4G* and *CLEC4M*) LSEC specific genes compared with day 12 iETV2-ECs (Fig. 3 A and S3A). Thus, iETV2-ECs could be maintained in HepMat hydrogels for 1 month, with stable tube formation for ±2 weeks.

**Figure 3.**
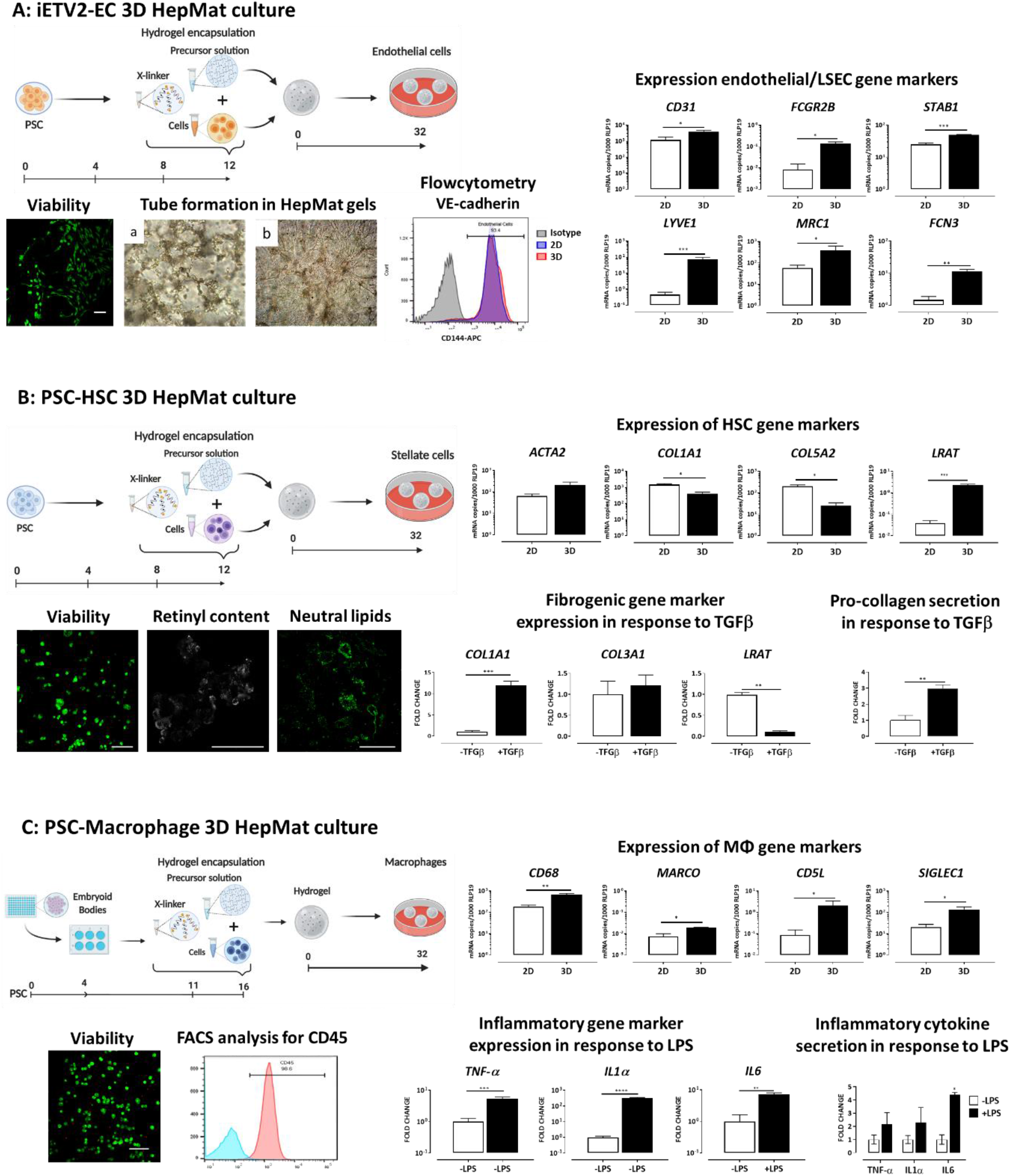
The HepMat hydrogel also maintains PSC non-parenchymal cells for at least 32 days. (A) PSC-endothelial cells (ECs): Schematic of culture of ECs from PSCs engineered to inducibly overexpress the EC master regulator *ETV2* (iETV2-ECs) initially for 12 days in 2D culture and then in HepMat hydrogels for 32 days. Live-dead staining with calcein-AM (green) and ethidium homodimer-1 (red) of d32 HepMat iETV2-ECs (representative for N=3 biological replicates). Bright field images of tube formation of iETV2-ECs cultured in HepMat hydrogel with (right panel) or without (left panel) ECGS. (Images taken on day 15 after embedding; representative for N=3 biological replicates). RT-qPCR for the general EC marker gene, *CD31*, and LSEC marker genes (*FCGR2B, STAB1, FCN3, LYVE1* and *MRC1*) in day 12 2D iETV2-progeny or recovered after an additional 32 days of culture in HepMat hydrogel (N=3 biological replicates). (iv) FACS analysis for VE-cadherin of iETV2-ECs cultured in 2D for 12 days (blue) vs. cultured for an additional 32 days in HepMat hydrogel (red) vs. isotype control (grey)(representative for N=3 biological replicates). **(B) PSC-hepatic stellate cells (HSCs):** Schematic of culture of HSCs from PSCs initially for 12 days in 2D culture and then in HepMAT hydrogels for 32 days. Live–dead staining with calcein-AM (green) and ethidium homodimer-1 (red) of PSC-HSCs after 32 days of culture in HepMat hydrogel (representative for N=3 biological replicates). UV-elicited autofluorescence typical for retinol A and BODIPY® staining of PSC-HSCs after 32 days of culture in HepMat hydrogel (representative for N=3 biological replicates). RT-qPCR for HSC marker genes (*ACTA2, COL1A1, COL5A2 and LRAT*) in day 12 2D HSC-progeny or recovered after an additional 32 days of culture in HepMat hydrogel (N=3 biological replicates). Treatment of PSC-HSC cultured in HepMat hydrogels for 32 days with 25ng/mL TGFβ for an additional 24h: Relative expression for HSC marker genes (*COL1A1, COL3A1, LRAT*) and pro-collagen secretion (by ELISA) in TGFβ-treated and non-treated cells (N=3 biological replicates). **(C) PSC-macrophages (Mϕs):** Schematic of culture of Mϕ from PSCs initially for +16 days in 2D culture and then in HepMAT hydrogels for 32 days. Live-dead staining with calcein-AM (green) and ethidium homodimer-1 (red) of PSC-Mϕ after 32 days of culture in HepMat hydrogel (representative for N=3). FACS analysis for CD45 on PSC-Mϕ cultured for 32 days in HepMat hydrogels (red) vs. isotype control (blue) culture (representative for N=3). RT-qPCR for Mϕ/KC marker genes (*CD68, MARCO, CD5L, SIGLEC1*) in day 12 2D Mϕ-progeny or recovered after an additional 32 days of culture in HepMat hydrogel (N=3 biological replicates). Treatment of Mϕ embedded in HepMat hydrogels for 32 days with 100 ng/mL LPS for an additional 24h: Relative inflammatory marker gene expression (*TNF-α, IL1α, IL6*) and TNF-α, IL1α, IL6 secretion (by ELISA) in LPS-treated and non-treated cells (N=3 biological replicates). Data are shown as mean ± SEM and analyzed by two-tailed student t-test. *p <0.05; **p <0.01; ***p <0.001; ****p <0.0001. Scale bars =100 µm.

We next differentiated PSCs towards HSCs using a previously published protocol (Coll et al., 2018). Day 12 HSCs expressed *ACTA2, PDGFRα, COL1A1, COL3A1, COL5A2* and *LOXL2* but low levels of *LRAT* (important for Vitamin-A metabolism in non-fibrotic HSCs) and more recently identified putative non-fibrotic HSC marker genes, *RGS5* and *IGFBP5* (Ramachandran et al., 2019)(Fig. 3 B and Fig. S3 B). Day 12 PSC-HSCs were then embedded in HepMat hydrogel, and maintained for 32 days in HSC medium. HSCs remained viable until day 32 in the HepMat hydrogel (Calcein-AM staining; Fig. 3 B). RT-qPCR on day 32 demonstrated a significant decrease in expression levels of *COL3A1* and *COL5A2*, and a significant increase in *LRAT* compared with HSCs harvested from 12-day 2D cultures, even if expression levels of other fibrotic (*COL1A1, ACTA2, LOXL2)* and non-fibrotic (*RGS5* and *IGFBP5)* genes remained unchanged (Fig.3B and S3 B). This transcriptional pattern suggests that culture of PSC-HSCs in 3D may induce at least a partial deactivation of HSCs (El Taghdouini et al., 2015; Leite et al., 2016; Mannaerts et al., 2015). Consistently, HSCs cultured in HepMat hydrogels stored retinol, as shown by BODIPY® staining and presence of typical blue autofluorescence elicited by UV light, even when cultured with low concentrations of palmitic acid (Fig. 3 B). To demonstrate functionality of the HepMat-embedded HSCs, we treated the cultures on d32 with 25 ng/mL TGFβ for 24 hours. RT-qPCR analysis demonstrated a significant increase in expression of *COL1A1* and a significant decrease in *LRAT* transcript levels (Fig. 3 B). TGFβ also induced significantly higher levels of secreted pro-collagen (Fig. 3 B). Thus, PSC-HSCs can be maintained for at least a month in HepMat hydrogels. Moreover, culture of HSCs in these hydrogels may at least in part induce a less fibrotic HSC phenotype while still allowing TGFβ-mediated activation.

Finally, we generated Mϕs from PSCs, using a protocol adapted from (van Wilgenburg et al., 2013) (Fig. 3 C). Day 16 PSC-progeny expressed typical Mϕ genes, including *CD163* and *CD45* but low levels of putative marker genes for liver specific Mϕs (i.e. KCs) such as *MARCO, CD5L* and *SIGLEC1* (Ramachandran et al., 2019). Day 16-PSC-Mϕs were then embedded in HepMat hydrogel, and maintained in macrophage medium. PSC-Mϕs remained viable until day 32 in HepMat hydrogels (Calcein-AM staining and FACS, Fig. 3 C). FACS demonstrated maintenance of CD45 expression on nearly 100% of cells (Fig. 3 C). RT-qPCR analysis demonstrated that typical Mϕ marker genes remained expressed. Interestingly, we found an increase of the putative KC marker genes, *CD68, MARCO* and *CD5L* (Fig. 3 C). To demonstrate functionality of the PSC-Mϕs after 32 days in HepMat cultures, we treated the cultures with 100 ng/ml lipopolysaccharide (LPS) for an additional 48 hours. LPS significantly induced expression of *TNF-α, IL1α and IL6* transcripts (Fig. 3 C) and an increased secretion of all three cytokines, which was significant for IL6 (Fig. 3 C). Thus, PSC-Mϕs could also be functionally maintained for at one month in HepMat hydrogels.

### Co-culture of HC3x-HLCs, iETV2-ECs, PSC-HSCs and PSC-Mϕs in hepatocyte maturation (HepMat) hydrogels

As the HepMat hydrogel, optimized to support HC3x-HLC progeny, also supported iETV2-ECs, PSC-HSCs and PSC-Mϕs, we next tested if the hydrogel would support co-culture of the four cell types, and if this would enhance the maturation/function of the different cells (Fig. 4 A). We first optimized different cell ratios and the medium composition. A ratio of HC3x-HLCs:iETV2-ECs:PSC-HSCs:PSC-Mϕs of 2:1:1:0.5 was optimal to retain all 4 cell types until d32 of co-culture (total number embedded 3×10^5^ cell/10μL). Supplementation of medium with AAGly, although beneficial for HC3x-HLC maturation (Fig. 2 A and S2 A), was toxic for iETV2-ECs (data not shown). Hence, the AAGly supplement was removed from the co-culture medium. When we did not add hematopoiesis supportive medium PSC-Mϕ did not survive in the co-culture system, the co-culture medium consisted of a 1:1 mix of LDM and StemPro™-34 SFM. Finally, all soluble ligands used in mono-cultures were incorporated in the final co-culture medium.

**Figure 4.**
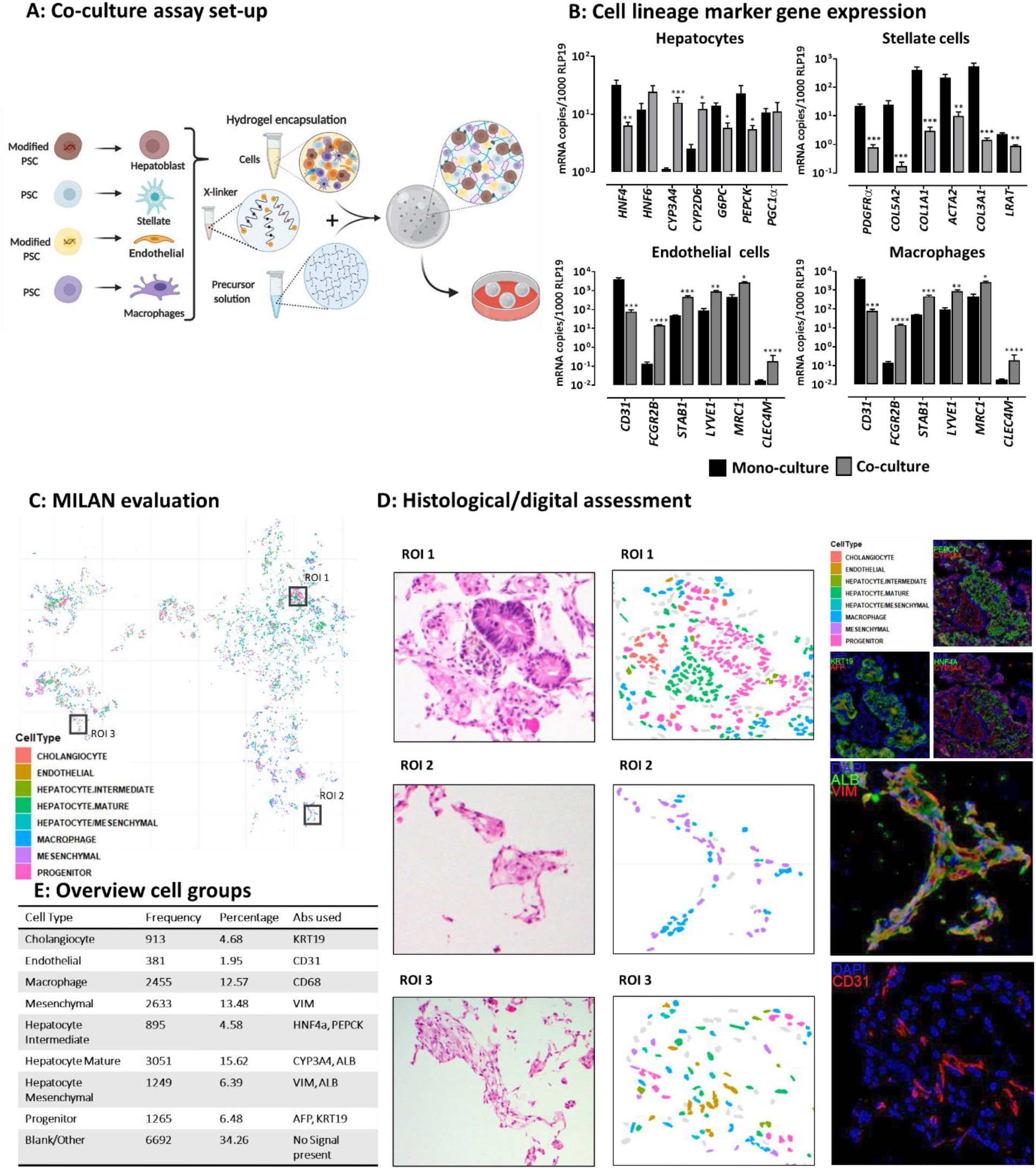
The HepMat hydrogel maintains and supports maturation /quiescence of PSC-hepatocyte-like cells, -endothelial cells, -hepatic stellate cells and -macrophages for at least 32 days. **(A)** Schematic representation of culture of HC3x-HLCs, iETV2-ECs, PSC-HSCs and PSC-Mϕs initially for 8-16 days in 2D culture and then in HepMat hydrogels for an additional 32 days. **(B)** RT-qPCR for marker genes for the different cell types: *HNF6, HNF4a, CYP3A4, CYP2D6, G6PC, PEPCK and PGC1α* as HLC markers; *CD31, FCGR2B, STAB1, LYVE1, MRC1 and CLEG4M* as general EC or LSEC markers, *PDGFRα, COL5A2, COL1A1, COL3A1, ACTA2* and *LRAT* as PSC-HSC markers and *MARCO, CD5L, SIGLEC1* and *CD68* PSC-Mϕ/KC markers in cells harvested after 32 days from respective 3D mono-cultures (3D) or 3D co-cultures (co-culture)(N=3 biological replicates (mono-culture data used in this figure are the same as in Fig. S2 and Fig.3)). **(C-E)** Multiple Iterative Labeling by Antibody Neodeposition (MILAN) analysis of 32 day HepMat hydrogels seeded with a combination of HC3x-hepatoblasts, ETV2-ECs, PSC-HSCs and PSC-Mϕs. Hydrogels were fixed without collagenase degradation, paraffin embedded and sectioned: example of digital reconstructed image after image processing of ALB, HNF4α, CYP3A4, PEPCK, AFP, CK19, CD31, CD68 and VIM (representative example of N= 4 independent hydrogels)(C). Digital reconstruction (middle) of regions of interest (ROI) with consecutive slices stained using HE (left) or immunofluorescent antibodies (right) staining. ROI1: ALB/CYP3A4+ and/or KRT19/AFP+, and/or PEPCK/HNF4α+; ROI2: ALB/VIM+ cells; ROI1: CD31+ cells (D); Overview of all cell types present obtained with the MILAN analysis after processing of the immunofluorescence images, unsupervised and supervised annotation of the clusters (E). Data are shown as mean ± SEM and analyzed by two-tailed student t-test.*p <0.05; **p <0.01; ***p <0.001.

We first analyzed marker gene expression for the different cell populations in the HepMat co-culture progeny. It should be kept in mind that this approach is compromised by the fact that the housekeeping gene transcript level is derived from all four cell types, and that some transcripts can be present in two or more cell populations (Aizarani et al., 2019). As the culture medium was no longer supplemented with the AAGly cocktail, we compared hepatic gene expression levels in d32 HepMat co-cultures with d32 HC3x-HepMat mono-cultures maintained in LDM without AAGly supplementation (Fig. S2 A). Transcripts for *CYP3A4* and *CYP2D6*, were significantly higher in HepMat co-cultures than HC3x-HepMat mono-cultures (Fig. 4 B). Consistently, BFC metabolization was significantly higher in HepMat co-cultures than in 3D-HepMat-HC3x mono-cultures, now reaching levels similar to suspension cultured PHHs (81±3.6%) (Fig S4 A). Again, CYP3A4 activity could be significantly induced by Rifampicin. This therefore suggests that co-culture of NPCs with HC3x-HLCs induces significant maturation of HC3x-HLCs, and this independent of AAGly required to induce maturation in 2D or 3D HLC-mono-cultures. Persistent expression of *CD31* (a typical large vessel EC marker; ±50-fold lower levels than in d12 2D iETV2-ECs) demonstrated that iETV2-ECs persisted until day 32 of co-culture (Fig. 4 B). Interestingly, a significant induction of *FCGR2B, STAB1, LYVE1* and *MRC1* was observed in HepMat co-cultures compared with iETV2-EC-HepMat mono-cultures, even if other LSEC marker genes were not increased (Fig. 4B and S4 B). Compared to PSC-HSC HepMat mono-cultures, fibrotic HSC transcripts decreased further in HepMat co-cultures, with 10-100 fold lower levels of *COL1A1, COL3A1, COL5A2, ACTA2* and *LOXL2*. The 2.5-fold reduction in *LRAT* levels, might be due to the dilution of HSCs in the mix of other cells (Fig. 4B). However, expression of *RGS5* and *IGFBP5* remained low (Fig. S4 B). Finally, expression of the Mϕ marker genes remained relatively unchanged in HepMat co-cultures compared to PSC-Mϕ HepMat mono-culture, even if *MARCO* expression levels were significantly higher in co-cultures than mono-cultures (Fig. 4 B and S4 B).

To gain a better understanding of the cellular composition of the co-culture system and to assess expression of each cell type separately rather than on a mixture of cells, H&E and cyclic multiplex immunofluorescence staining (Bolognesi et al., 2017; Bosisio et al., 2020) were performed. HepMat co-cultures were fixed and embedded on day 32 (Fig. S4 C). Histological and Multiple Iterative Labeling by Antibody Neodeposition (MILAN) analysis of d32 HepMat co-cultures revealed continued presence of progeny from the four different embedded cell types (Fig. 4 C & D; and Fig. S4 E&F). The co-cultures showed characteristics of one coherent functional unit with a balanced distribution of each of the cell types. The epithelial compartment comprised hepatocyte-like cells at different stages of development: AFP-/ALB+/CYP3A4+ mature hepatocytes, AFP+/ALB+/CYP3A4-intermediate hepatocytes, as well as a fraction of AFP+/KRT19+ hepatic progenitors. We also detected VIM+/ALB+ mesenchymal hepatocytes. Interestingly we also could detect ductular structures that stained were AFP-/KRT19+, a marker profile for cholangiocytes, consistent with the remaining bipotency of the hepatoblasts harvested on day 8 from 2D cultures used to initiate the culture (Fig. S4 D). Immunofluorescence stains also revealed presence of flat CD31+ ECs that were lining the epithelial components. In addition, CD68+ Mϕs and VIM+ mesenchymal cells were scattered between these structural components. This was confirmed by further sectioning of additional hydrogels and H&E staining as well as immunostaining, analyzed by confocal microscopy as shown in Fig. S4 G.

Thus, the HepMat co-culture system supported maturation of PSC-hepatoblasts to cells with a mature hepatocyte like phenotype (as well as a fraction of cells with cholangiocyte features), induced less fibrogenic PSC-HSCs and an apparent more LSEC committed EC progeny.

### HepMat HC3x-HLC, iETV2-EC, PSC-HSC and PSC-Mϕ co-cultures can be used to model liver steatosis and fibrosis

In a final set of studies, we addressed if HepMat co-cultures could be used to model liver fibrosis, steatosis and inflammation, by exposing cultures either to TGFβ or oleic acid (OA). We treated d32 HepMat co-cultures with 25 ng/ml of TGFβ for an additional 3 days (single administration) or 7 days (2 administrations). In HepMat co-cultures, exposure to TGFβ caused a significant increase in transcripts for *COL1A1* (17±3.5-fold) and *COL3A1* (13±3.4-fold), but not *COL5A2* (Fig. 5 A). TGFβ exposure for 7 days also induced a 7±0.49 and 13±4.3-fold induction of *IL1α* and *IL6* transcripts, respectively (Fig. 5 A). Comparatively, when mono-cultures of HC3x-HLCs, PSC-HSCs or PSC-Mϕs were exposed to a similar schedule of TGFβ, a significant but much smaller induction of *COL1A1* transcripts was observed in HC3x-HLC (9±1.6-fold 3 days treatment; 5±0.7-fold 7 days treatment) and *COL1A1* and *COL3A1* transcripts in PSC-HSC (4±1.1-fold d3). TGFβ did not induce inflammatory gene expression in PSC-HLC mono-cultures and only a modest increase in PSC-HLC or PSC-Mϕ monocultures. In mono-culture, iETV2-ECs did not survive the TGFβ treatment; therefore, no data is shown. In addition, TGFβ induced a 7±0.4-fold higher secretion of pro-collagen in HepMat co-culture supernatants on day 7, and a 17±3.4-fold higher level of IL6 (Fig. 5 B). Pro-collagen levels were unchanged in HC3x-HLCs and PSC-Mϕ mono-cultures, and ±3-fold increase in PSC-HSC monocultures 3 and 7 days after treatment with TGFβ (Fig. 5 B). Hence, fibrogenic responses were more robust when an interplay between NPCs and hepatocytes was possible in HepMat co-cultures compared with either of the HepMat mono-cultures.

**Figure 5.**
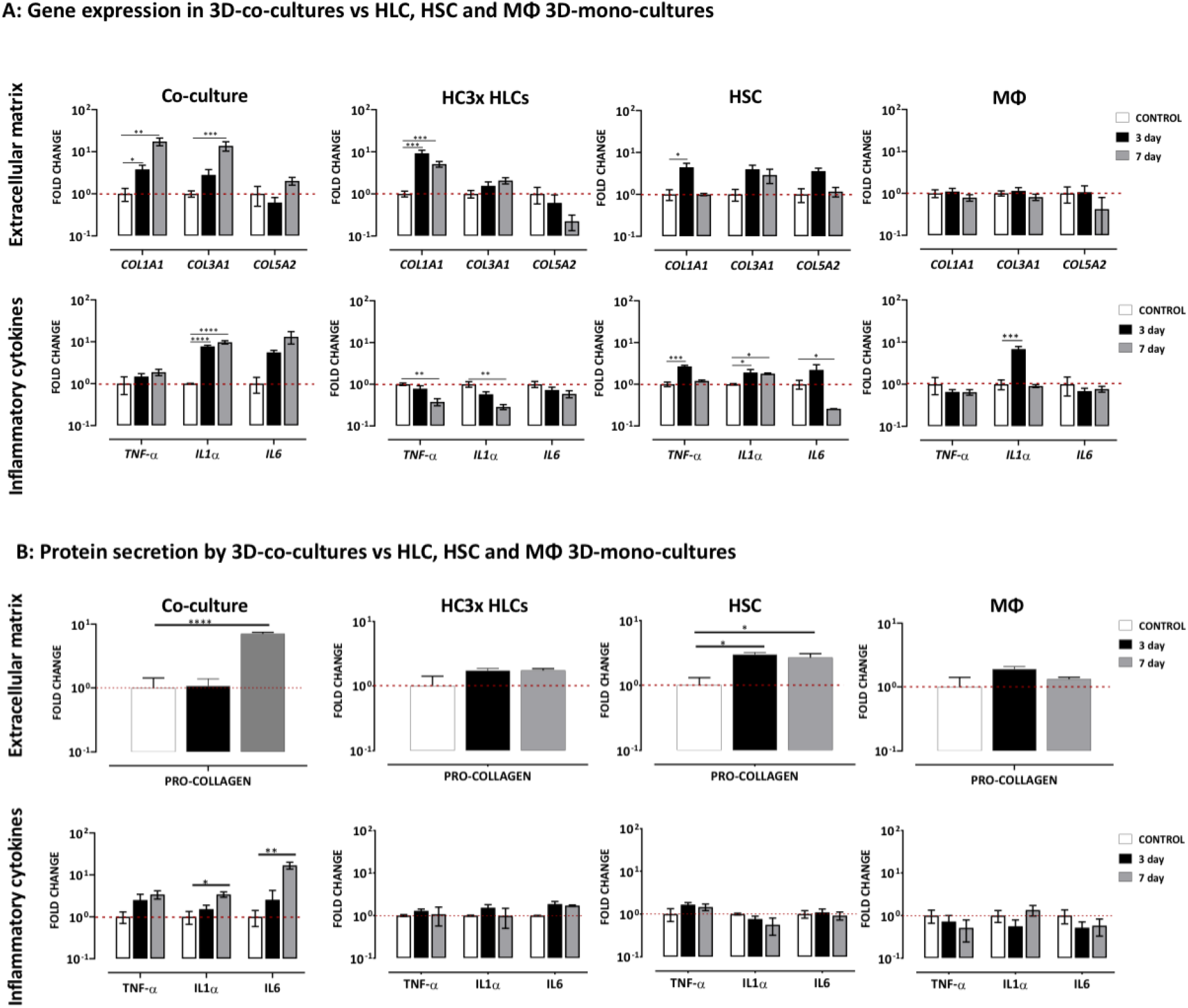
HepMat-based co-cultures of PSC-hepatocyte-like cells, -endothelial cells, -hepatic stellate cells and -macrophages can be used to model TGFβ-induced liver fibrosis. **(A)** Relative expression of fibrogenic markers (*COL1A1, COL3A1* and *COL5A2*) and pro-inflammatory cytokines (*TNF-α, IL1α* and *IL6*) in d32 HepMat 4-cell co-cultures or PSC-HC3x-HLC, PSC-HSC or PSC-Mϕ monocultures exposed to TGFβ for 3 and 7 days (N=3 biological replicates)(red horizontal line represents the control sample). **(B)** Relative measurement of secreted protein pro-collagen and pro-inflammatory cytokines (TNF-α, IL1α and IL6) in d32 HepMat 4-cell co-cultures or PSC-HC3x-HLCs, PSC-HSCs or PSC-Mϕ mono-cultures exposed to TGFβ for 3 and 7 days (N=3 biological replicates) (red horizontal line represents the control sample). Data are the mean ± SEM (N = 3) and analyzed by one-way Anova (Tukey’s multiple comparison).*p <0.05; **p <0.01; ***p <0.001, ****p <0.0001.

We also exposed d32 HepMat co-cultures to 0.8 mM OA for 3 days (1 dose) and 7 days (2 doses) to evaluate steatosis and fibrosis induction (Ouchi et al., 2019). BODIPY® staining revealed significant lipid accumulation in co-cultures treated with OA already on day 3, which persisted until day 7 (Fig. 6 A). In HepMat co-cultures, OA induced a 14±2.3-to 18±1.3-fold induction of *COL1A1* and *COL3A1* already after 3 days, and an even greater induction of *COL1A1, COL3A1* and *COL5A2* on day 7 (Fig. 6 B). Moreover, OA also massively induced *IL6* expression (1200-fold d3; 1000-fold d7) and induced a 5±2.3-to 19±3.4 -fold induction of *TNF-α* and *IL1α* on day 3 (Fig. 6 B). These responses were much more modest in mono-cultures, with a ±9-fold induction in *COL1A1* and *COL3A1* transcripts 3 days after OA addition, and a significantly lower induction of inflammatory cytokines in HC3x-HLC and PSC-Mϕ mono-cultures. As iETV2-ECs did not survive OA treatment in mono-cultures, no data is shown. Consistent with the RT-qPCR data, a 5±0.5 and 12±1.5 -fold increase in pro-collagen was observed in supernatants of 3 and 7 day HepMat co-cultures after administration of OA, and a 90±5-fold increase in IL6 on day 7. By contrast, OA induced no significant changes in either pro-collagen or inflammatory cytokines in supernatants of any of the mono-cultures (Fig. 6 C). Thus, the significantly greater induction of both fibrogenic genes and inflammatory cytokine genes (especially *IL6*) in OA treated HepMat co-cultures compared with the different mono-cultures is consistent with the notion that liver inflammation and fibrosis in response to lipids (as in NASH) requires an interplay between NPCs and hepatocytes.

**Figure 6.**
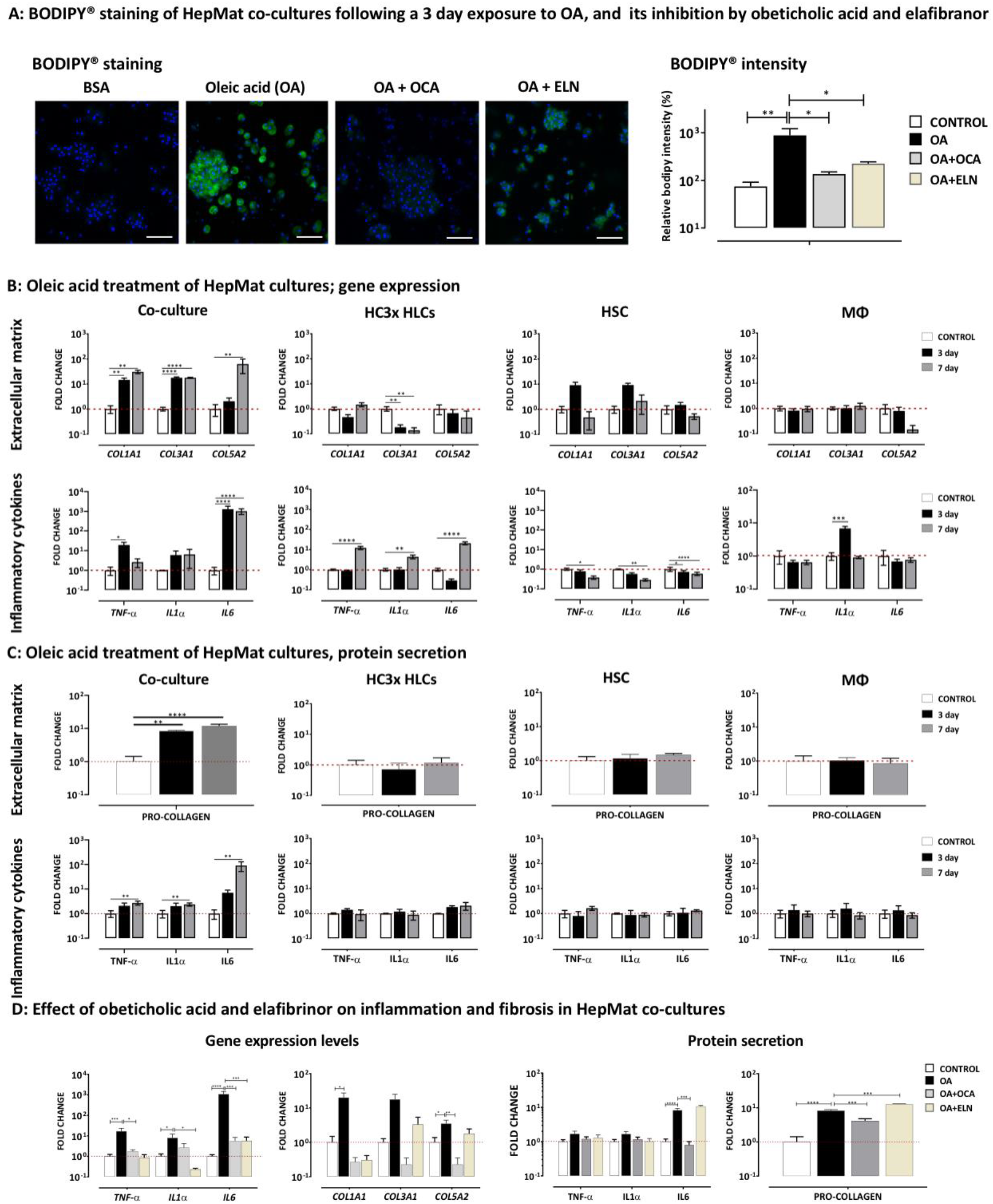
HepMat-based co-cultures of PSC-hepatocyte-like cells, -endothelial cells, -hepatic stellate cells and -macrophages can be used to model liver steatosis, fibrosis and inflammation. **(A)** Representative confocal fluorescence images of d32 HepMat co-cultures exposed for 3 days to BSA as control, oleic acid (OA; 800µM), a combination of oleic acid + obetecholic acid (OA; 800µM +OCA; 1μM) or oleic acid + elafibranor (OA; 800µM+ELN; 30 µM); (N=3 biological replicates)(scale bar = 100 µm); Relative BODIPY® intensity compared with BSA control on day 3 and day 7 (N=3 biological replicates). **(B)** Relative expression of fibrogenic markers (*COL1A1, COL3A1* and *COL5A2*) and pro-inflammatory cytokines (*TNF-α, IL1α*, and *IL6*) in d32 HepMat 4-cell co-cultures or PSC-HC3x-HLCs, PSC-HSCs or PSC-Mϕ monocultures exposed to OA for 3 or 7 days (N=3 biological replicates)(red horizontal line represents the control sample; control sample data used in this figure are the same as in Fig. 5) **(C)** Relative measurement of secreted pro-collagen and pro-inflammatory cytokines (TNF-α, IL1α and IL6) in d32 HepMat 4-cell co-cultures or PSC-HC3x-HLCs, PSC-HSCs or PSC-Mϕ mono-cultures exposed to OA for 3 or 7 days (N=3 biological replicates)(red horizontal line represents the control sample). **(D)** Relative expression of fibrogenic and proinflammatory marker genes, and secreted pro-collagen, TNF-α, IL1*α* and IL6 in d32 HepMat 4-cell co-cultures exposed to BSA, OA alone, OA & OCA or OA & ELN for 3 days (N=3 biological replicates)(red horizontal line represents the control sample). Data are the mean ± SEM (N = 3) and analyzed by one-way Anova (Tukey’s multiple comparison).*p <0.05; **p <0.01; and ***p <0.001, ****p <0.0001. OA: Oleic acid; OCA: Obeticholic acid; ELN: Elafibranor.

In a final set of studies, we tested the effect of 2 late phase III anti-NASH drugs, namely the farnesoid X receptor (FXR) agonist obeticholic acid (OCA), which has shown improvement in fibrosis in 23% of patients vs. placebo treated patients (Younossi et al., 2019), and the PPARα/δ activator, elafibranor (ELN), which was recently (https://ml-eu.globenewswire.com/Resource/Download/38e085e1-66f5-4251-8abe-648d0e7b9ed1) shown to not significantly improve liver fibrosis compared to placebo-treated patients. When HepMat co-cultures were co-treated with either OA & OCA or OA & ELN for 3 days, a significant reduction in BODIPY® staining was observed (Fig. 6 A). OCA robustly decreased *COL1A1* and *COL3A1* levels, while ELN only inhibited *COL1A1* transcripts; which was translated in a significant albeit not complete inhibition of pro-collagen secretion by OCA and no effect from ELN on pro-collagen concentrations in culture supernatants (Fig. 6 D). A similar effect was observed on proinflammatory parameters (Fig. 6 D). Both OCA and ELN significantly reduced *IL6* transcripts, but only OCA could inhibit production of secreted IL6.

## Discussion

We here describe the creation of a fully tunable and synthetic PEG-based hydrogel, functionalized with a combination of ECM and CAM binding sequences (termed hepatocyte maturation hydrogel, or HepMat hydrogel) that supports the maintenance and maturation of PSC-HLCs progeny. In line with our hypothesis that such a hydrogel composition should also allow maintenance of NPCs, co-localized *in vivo* with hepatocytes in liver sinusoids, we demonstrated that HSCs and Mϕs generated from PSCs survive in the HepMat hydrogel for at least 32 days, while maintaining their fibrogenic response to TGFβ and inflammatory response to LPS. PSC-ECs (generated by overexpression of the master regulator *ETV2* (Elcheva et al., 2014), could also be maintained for at least 32 days. Co-culture of PSC-HLCs and PSC-NPCs in a 4-cell type HepMat-co-culture system further enhanced HLC maturation, supported persistent vessel-like structure establishment and possible fating of ECs to cells with a more LSEC-like phenotype, induced a less activated state of HSCs, and possibly induced some KC like features in Mϕs. Most importantly, the 4-cell co-culture had a far greater fibrogenic and inflammatory response to TGFβ and, even more so, OA, than any of the mono-cultures. This attests to the requirement of an intricate interaction between NPCs and HLCs in the development of liver fibrosis and inflammation, and the capability of the HepMat-PSC-HLC-NPC co-culture to model liver fibrosis/inflammation. As we could block the pro-inflammatory and -fibrogenic effects of OA by treatment of the 4 cell-type HepMat co-culture with obeticholic acid, this culture system should be very helpful in developing anti-NASH / fibrogenic drugs.

It has been long-established that when cultured on stiff culture plates, PHHs very quickly lose functionality (Boon et al., 2020; Godoy et al., 2016). This is also the case for HSCs that become quickly activated (El Taghdouini et al., 2015; Mannaerts et al., 2015), and LSECs that very quickly lose their typical sinusoidal endothelial characteristics (Sorensen et al., 2015). Especially for hepatocytes, numerous studies have demonstrated that this can at least in part be prevented when cells are cultured for instance in sandwich cultures, in spheroids or embedded in scaffolds (Bell et al., 2018; Godoy et al., 2016; Hurrell et al., 2020; Jaramillo et al., 2018; Ma et al., 2016; Mazzocchi et al., 2018; Nakai et al., 2019; Prestwich, 2011; Toivonen et al., 2016).

In initial studies, we tested the maturation of HLCs in different natural polymers, by embedding day 8 hepatoblasts in collagen, matrigel, and or gelatin gels for up to 28 days. Although culture in a mixture of gelatin and matrigel increased some hepatic marker transcripts, the hydrogels were rapidly degraded and hence did not improved HLC maturation (Fig. S5A & B). We therefore created PEG-based hydrogels crosslinked in a non-cytotoxic manner with linkers that can be cleaved by MMPs thought to be present in liver (Duarte et al., 2015; Naim et al., 2017). We chose linkers that would be relatively slowly degraded (Patterson et al., 2010) to ensure stability of the hydrogel for at least 30 days. As PEG is inert and does not allow cell adhesion, the PEG gel can be mixed with natural polymers. However, batch-to-batch variability of the natural polymers may prevent creation of highly consistent culture conditions. We therefore decorated the PEG hydrogel with peptide ligands representing cell adhesion domains of ECM molecules and or CAMs, present in human livers (Naba et al., 2016).

We used a DOE approach to enable screening the innumerable combinations of hydrogels, based on PEG concentration and degradability by MMP-cleavage and the 24 different peptides selected. The DOE narrowed down the number of combinations to be tested to 216 different environments, whereas a non-DOE approach would have required screening of >20,000 conditions. We found that four-arm PEG with stiffness of 9Kpa, crosslinked with the MMP9 cleavable linker peptide, GCRD<u>GPQGIAGQ</u>DRCG, and functionalized by a fibronectin [P5], laminin [P8], collagen I [P12 and P13], collagen IV [P16] and E-cadherin [P23] peptide (Table 1) generated PSC-HLC that aligned transcriptionally and functionally significantly closer to PHHs than 2D cultured PSC-HLCs. Interestingly, some hydrogel compositions had a detrimental effect on HLC maturation compared to 2D culture (Fig. 1 and Fig. S1). Future studies focusing on these hydrogels might be of interest to further understand how HLC maturation occurs *in vitro*, and how this might be further enhanced, by counteracting mechanisms operative in these non-supportive gels.

We recently demonstrated that significantly improved CYP450 activity, drug biotransformation and cellular metabolism of PSC-HLCs can be obtained when three hepatic transcription factors (HC3x) are inducibly overexpressed during differentiation and cells are allowed to mature in amino acid rich (AAGly) medium, even in 2D cultures (Boon et al., 2020). We here demonstrate that when such HC3x progeny is maintained in 3D HepMat cultures and culture in with AAGly supplemented medium, a significant further transcriptional and functional maturation can be obtained. We observed a significant induction of genes involved in gluconeogenesis and mitochondrial biogenesis and significantly enhanced and rifampicin-inducible CYP3A4 activity. Interestingly, when HC3x-HLCs were maintained in HepMat hydrogels in baseline LDM medium not supplemented with extra AAs, HLC functionality remained significantly inferior to the HepMat-AAGly culture condition, demonstrating the importance of both the functionalized hydrogel and AA support for HLC maturation, in HLC mono-cultures.

The HepMat formulation could also maintain PSC-HSC (as described in (Coll et al., 2018)) for at least one month. Interestingly, culture of PSC-HSCs in the HepMat hydrogel induced a transcriptional profile more reminiscent of non-fibrotic HSCs, even if not all the recently described (Ramachandran et al., 2019) non-fibrotic HSC gene markers were induced. This is consistent with the notion that culture of HSCs in 3D may counteract activation observed when cells are cultured on stiff surfaces (Mannaerts et al., 2015; van Grunsven, 2017). Noteworthy, a fibrogenic response of HSCs, embedded in the HepMat hydrogel was still feasible, as shown by incubation with TGFβ. This gene induction far exceeded what we previously reported for TGFβ treated PSC-HSCs (or primary human HSCs) (Coll et al., 2018; Leite et al., 2016), likely because PSC-HSCs display a significantly less fibrotic phenotype in HepMat 3D cultures compared with 2D cultured cells. This indicates that PSC-HSCs cultured in the 3D-HepMat hydrogel might be a good model, in itself, to study HSC activation and examine drugs that might counteract fibrogenic responses. The HepMat hydrogel supported survival of PSC-ECs for 32 days, even if spontaneous tube formation ceased after 15 days. This is likely due to the absence of cells, such as mural cells, known to stabilize EC tube formation *in vivo* and *in vitro* in these mono-cultures (Gaengel et al., 2009). Finally, HepMat hydrogels also supported the survival of PSC-Mϕs (van Wilgenburg et al., 2013) for at least 32 days beyond the 16+ days of culture in 2D suspension differentiation cultures, and, even following 32 days in hydrogel culture, PSC-Mϕs could be activated by LPS to produce inflammatory cytokines. As the ECM/CAM interaction requirements of some/all of the NPCs may differ in some aspects from that of hepatocytes, it may be of future interest to determine if the HepMat hydrogel might be further functionalized with additional ECM/CAM peptides for an even better support of the different NPCs.

We then co-incorporated all four PSC-derived cell populations in the HepMat gel to create an all-PSC derived 3D liver model, and test the model for its suitability to study liver inflammation and fibrosis. Similar to the spheroid-like clusters that were formed when single-cell suspensions of HC3x-HLCs were cultured in HepMat hydrogels, single cell suspensions of HC3x-HLCs co-embedded together with PSC-NPCs also led to the creation of spheroid-like structures. In addition, in the co-cultures we could also see the appearance of tubular structures interspersed between the spheroid structures. This apparent spontaneous morphogenesis the 4-cell co-culture model has not been fully characterized, but light microscopy demonstrated that this occurred during the first 10 days of (co)-culture. The observation that the input cell ratio (44% HC3x-hepatoblasts, 22% iETV2-ECs, 22% PSC-HSCs, 11% PSC-Mϕ), was similar to the composition of cells present in the 4-cell co-culture, except for an apparent loss of ECs (of the identified cell types, 57% were endodermal, 20% mesenchymal and hematopoietic but only 2% endothelial), argues against the fact that cell proliferation in situ underlies the creation of spheroids or tubular structures.

One could argue that the creation of multicellular structures within the hydrogel is in itself responsible for the maturation and further cell fating of HC3X-HLCs, and that presence of the HepMat hydrogel plays only a minor role. Arguments against this are, first, that removal of a single peptide from the peptide pool resulted in inferior HLC maturation (data not shown). Second, HLC maturation was highly dependent on different peptide combinations, different stiffness or degradability of the hydrogels, and some hydrogels even decreased maturation of HLCs compared to 2D cultures. Third, although we could generate HLC-spheroids by forced aggregation, significantly inferior maturation was observed compared with HLCs matured in HepMat hydrogels (data not shown). Finally, we have not yet succeeded in creating self-assembling spheroids from all four cell types together. This therefore strongly suggests an instructive effect of the HepMat hydrogel on cell maturation.

Multiplex immunostaining revealed that of all endodermal progeny in HepMat co-cultures, 10% had a phenotype of hepatic progenitors (AFP+/KRT19+) and 7% an intermediary (AFP+/ALB+/CYP3A4-) hepatocyte phenotype, while 44% of cells displayed an AFP-/ALB+/CYP3A4+ staining pattern, compatible with mature hepatocytes. In addition, ±18.4% of cells co-stained with hepatocyte and mesenchymal markers, suggesting presence of some degree of epithelial-to-mesenchymal transition (EMT). Finally, we also detected 13.4% of cells with a cholangiocyte phenotype, suggesting the creation of biliary ducts from the d8-PSC-hepatoblasts. Cholangiocyte differentiation from bipotent hepatoblasts is governed by a number of cell-extrinsic signals emanating from mesenchymal structures adjacent to bile ducts (reviewed in (Lemaigre, 2020)). This includes signaling by TGFβ derived from the periportal mesenchyme to commit hepatoblasts to ductal plate cells, and JAGGED1, also expressed in the periportal mesenchyme, which activates the NOTCH2 signaling pathway to support cholangiocyte differentiation. Therefore, the development of biliary ducts in co-cultures of PSC-hepatoblasts and NPCs may not be surprising. Vascular tube-like structures persisted until at least 32 days in the combinatorial system, which was in stark contrast with PSC-iETV2-EC mono-cultures, consistent with the need for supporting cells for stable vasculogenesis.

Not unexpectedly (Ardalani et al., 2019; Koui et al., 2017; Takebe et al., 2013; Wang et al., 2018), presence of PSC-NPCs supported HLC maturation. As AAGly, crucial for maturation of HC3x-HLC in 2D culture (Boon et al., 2020) and in HepMat mono-culture, appeared toxic for PSC-iETV2-ECs, we removed the AAGly supplement from the co-culture medium. Nevertheless, even when we omitted AAGly, *CYP450* gene expression as well as function reached levels similar to PHHs, demonstrating that the co-culture system in its own supports HLC maturation and AA-supplementation of the culture medium is no longer required.

In addition, co-culture of PSC-iETV2-ECs with the other cell populations induced LSEC-like gene expression, as we observed a significant further induction of *FCGR2B, LYVE1, MRC1* and *STAB1*. One word of caution is that even though these marker genes are commonly used to demonstrate presence of LSECs in primary hepatocyte/NPC spheroid cultures (Baze et al., 2018; Messner et al., 2013), promiscuous expression of scavenging marker genes in KC/Mϕs and LSECs exists (Aizarani et al., 2019; Ramachandran et al., 2019). Other newly identified LSEC marker genes (such as *FCN3, OIT3, CLEC4M, CLEC4G*) were not significantly higher in HepMat co-cultures compared with iETV2-EC HepMat mono-cultures, but this may in part be because the relatively limited number of iETV2-ECs remaining in the final mixture. In the co-culture, we also noted a further decrease in fibrotic HSC markers, including multiple collagen genes, *PDGFRα*, the quintessential fibrotic niche associated HSC marker (Ramachandran et al., 2019), as well as *ACTA2* and *LOXL2*. Nevertheless, other putative non-fibrotic HSC marker genes such as of *RGS5* and *IGFBP5*, identified by single cell RNA sequencing of non-fibrotic livers (Ramachandran et al., 2019), remained low in the HepMat co-cultures. PSC-Mϕ persisted, even though most putative KC-specific gene markers (Ramachandran et al., 2019) were not higher in the co-culture system compared to PSC-Mϕ-3D mono-cultures.

When the co-cultures were exposed to either TGFβ or OA, we found a massive increase in pro-fibrogenic and pro-inflammatory genes, compared with any of the 3D mono-cultures, demonstrating that liver inflammation and fibrosis is, as *in vivo*, dependent on an intricate interaction between HLCs and NPCs (Marrone et al., 2016). The induction of e.g. secreted IL6 in HepMat co-cultures appeared to exceed levels reported previously for PSC-multi-cellular human liver organoids (HLOs)(Ouchi et al., 2019). This might be due to the higher frequency of especially macrophages in HepMat co-cultures (11% in current study vs. 0.8% in HLOs). Moreover, the induction of collagens (e.g. *COL1A1* transcripts) by TGFβ exposure was similar as for liver spheroids consisting of PHHs and the most fibrogenesis-inducing primary liver derived NPCs (Hurrell et al., 2020). It will be of interest in future studies to perhaps compare co-culture of HC3x-HLCs with any of the three PSC-derived NPCs separately, to gain insight in how their combined presence induces this fibro/inflammatory response. The steatosis (evaluated by BODIPY® staining), pro-inflammatory and fibrogenesis inducing effect of OA could be largely blocked by treatment of the culture with OCA, a drug shown in phase III studies to block fibrosis and / or decrease NASH features (Younossi et al., 2019). By contrast, Elafibranor, which was recently shown to not improve liver fibrosis compared to placebo treatment in a phase III study, did not inhibit OA induced secretion of collagen and IL6, even if it decreased some of the fibrogenic and inflammatory gene transcript levels, and decreased steatosis, albeit somewhat less than following OCA treatment.

We believe that the approach to create functional multicellular liver models described here is likely generic. It would therefore be of interest to test similar approaches to create functional multicellular co-culture systems starting from pluripotent stem cells, to create for instance brain-like tissues containing both ectodermal and mesodermal descendants, among others. Furthermore, the notion of creating a fully defined tissue-mimicking hydrogel formulation, with tunable stiffness, degradability and organ-specific-ECM and -CAM peptide-based functionalization, could also aid in creating tissue-like cultures and organoids starting from tissue derived (stem) cells.

In conclusion, we created an all-PSC-derived hepatocyte- and NPC-like cell co-culture system that significantly improves PSC differentiation and specification to mature hepatic progeny, PSC-HSCs with a less fibrotic but very responsive phenotype, and PSC-ECs with a more LSEC phenotype. As the co-culture is based on a fully defined hydrogel composition and well-defined ratios of hepatoblast- and NPC input, the culture system may be less variable than spontaneous PSC-co-differentiation cultures with or without natural polymer matrices. Finally, the co-culture system is suitable for studying liver steatosis, inflammation and fibrosis as well as assess drugs counteracting these effects.

## Acknowledgements

This study was funded by grants to M.K. (European Union’s Horizon 2020 Research and Innovation programme under the Marie Sklodowska-Curie-IF grant agreement number 657701), J.D.S. (Fonds Wetenschappelijk Onderzoek 1S33916N), R.B. (Agentschap voor Innovatie door Wetenschap en Technologie SB-121393), T.T. (Fonds Wetenschappelijk Onderzoek 1185918N), J. DB. (Fonds Wetenschappelijk Onderzoek 1189120N), A.A. (KU Leuven funding (C3/19/053), Opening the Future Campaign of the KU Leuven University Fund), F.M.B. (KUL interne fondsen middel-zware infrastructuren EMH-D8191-AKUL/19/30 I005920N), C.M.V. and L.v.G. (IWT-140045, HILIM-3D). C.M.V. (Fonds Wetenschappelijk Onderzoek G0D9917N; also this project has received funding from the European Union’s Horizon 2020 research and innovation programme under grant agreement No 681002 (EU-ToxRisk)).

Confocal images were recorded on a Zeiss LSM 880 (Cell and Tissue Imaging Cluster (CIC), Supported by Hercules AKUL/15/37_GOH1816N and FWO G.0929.15 to Pieter Vanden Berghe, University of Leuven.

Stiffness measurements were performed at the Flanders Institute for Biomechanical Experimentation (FIBEr), supported by KU Leuven Research Fund (grant no. IDO/13/016 to C.M.V. and Dr. Hans Van Oosterwyck).

## Contributions

M.K. designed and planned the study, generated the data, and wrote the manuscript. B.T. helped with co-culture optimization, hydrogel characterization and histological analyses. M.V.H. and A.A. helped with multiplexing stains and MILAN analysis. R.B. created the HC3x and iETV2 PSC-lines. J.D.S. helped with the analysis using ‘R’. F.C. helped with the optimization of HepMat monocultures; T.T. and J.DB. helped with optimizing differentiations and P.T, M.C. and T.I. I. helped with optimization of the co-culture system. M.B. helped with FACS experiments and analysis. F.M.B. and T.R. provided expertise in 3D mono- and co-culture histological analysis, and MILAN analysis. A.R. helped with the analysis to define the optimal HepMat composition. L.v.G provided scientific input, helped with data interpretation, and contributed to writing the manuscript. C.M.V designed the study and helped with the analysis of the data and writing of the paper. All authors read and edited the manuscript.

## SUPPLIMENTARY FIGURES

**Figure S1.**
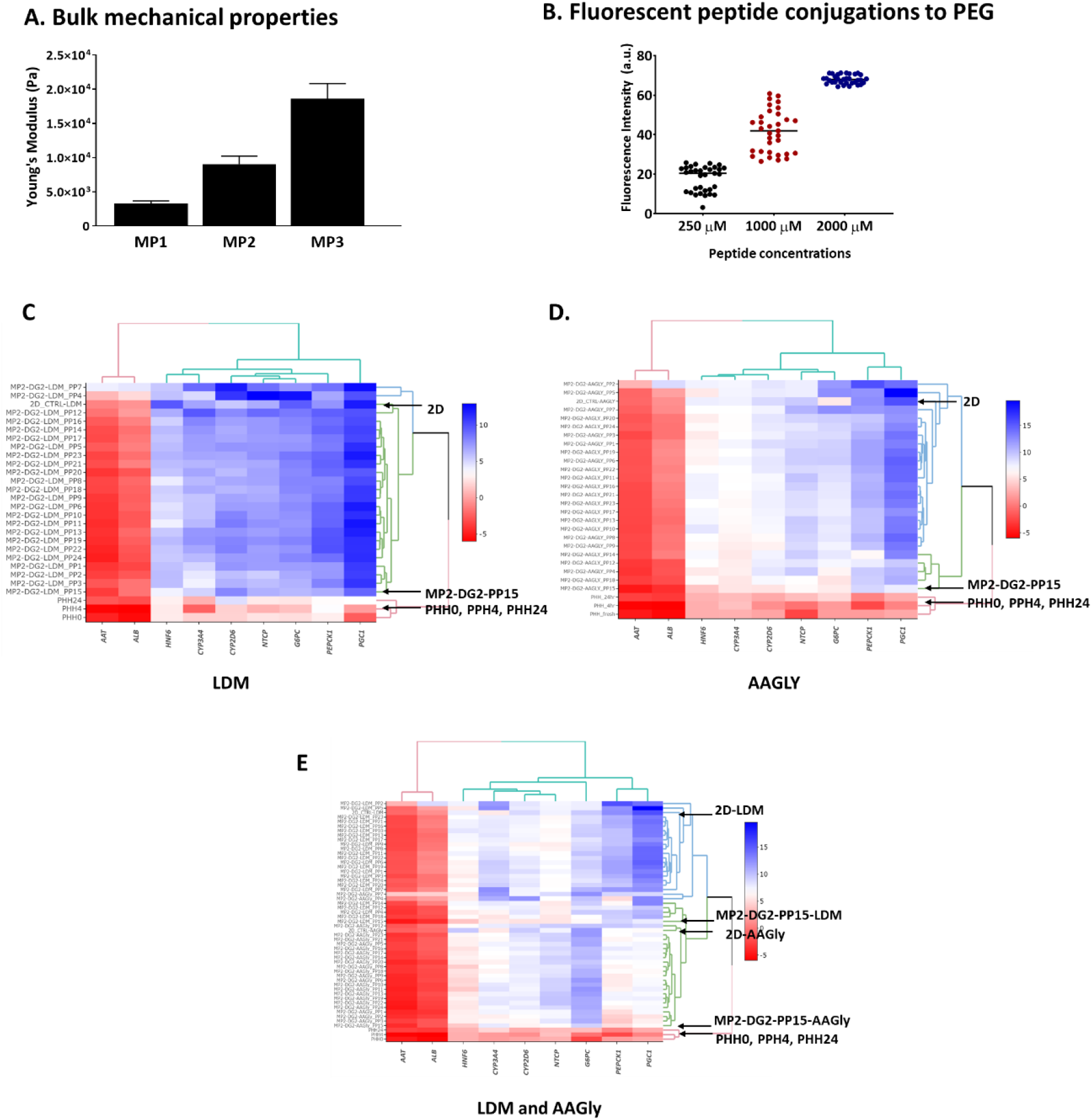
Confirmatory screens demonstrate that the MP2-DG2-PP15 hydrogel optimally supports HLC differentiation, both in basic liver differentiation medium (LDM) and amino-acid-glycine supplemented (AAGly) medium. **(A)** Bulk mechanical properties measured by nanoindentation of hydrogels containing 3 different concentrations of PEG polymers, crosslinked with DG2, and functionalized with 6 different functionalizing peptides (N=2 biological replicates). **(B)** Fluorescent intensity measurements obtained from fluorescent 5-FAM peptide conjugated PEG hydrogels at different concentrations. Each dot represents fluorescent intensity measured at different planes in the z-axis of hydrogels conjuf=gated with the 3 different 5-FAM concentrations (150, 1000 and 2500μM) (N= 1 biological replicates). **(C-E)** Hierarchical clustering analysis based on transcript levels of 8 hepatocyte marker genes in day 20 PSC-HLC progeny cultured in MP2-DG2 hydrogels functionalized with different peptide pools (PP1-P24) compared with PHH0, PHH4 and PHH24 and HLCs in 2D culture in basic LDM medium (C), AAGly medium (as described in Boon et al., 2020)(D), and a combined LDM and AAGly analysis (E); compared with fresh PHH0, PHH4 and PHH24. (Note: Cluster gram shows delta CT compared with the *RPL19* housekeeping transcript; red is higher expression and blue lower expression). MP = mechanical property based on PEG concentration; DG: degradation property, based on MMP cleavable linker; PHH = primary human hepatocytes; LDM = liver differentiation medium; AAGly medium is LDM + extra amino acids.

**Figure S2.**
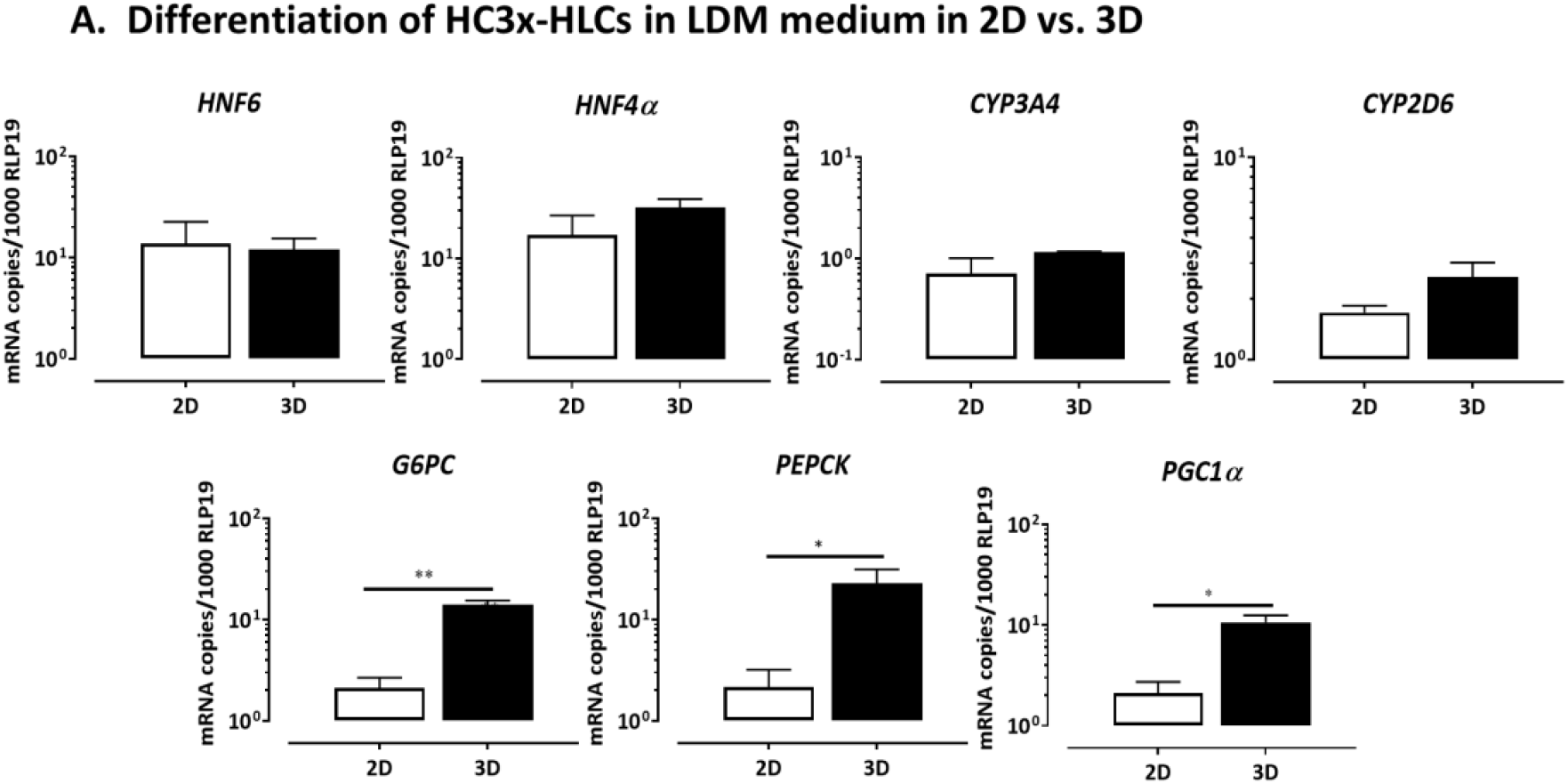
The HepMat hydrogel supports further maturation of PSC-HLC. **(A)** HC3x-PSCs were cultured either for 40 days in 2D culture or initially for 8 days in 2D culture and then in HepMat hydrogels for 32 days using liver differentiation medium without extra amino acids (Boon et al., 2020). Gene expression of *HNF4α, HNF6, CYP3A4, CYP2D6, G6PC, PEPCK and PGC1α* in day 40 HC3x-HLCs maintained in 2D culture vs. progeny from HepMat hydrogel cultures (N=3 biological replicates). Data are shown as mean ± SEM and analyzed by two-tailed student t-test; *p <0.05; **p <0.01Figure 2. The HepMat hydrogel supports further maturation of PSC-HLC.

**Figure S3.**
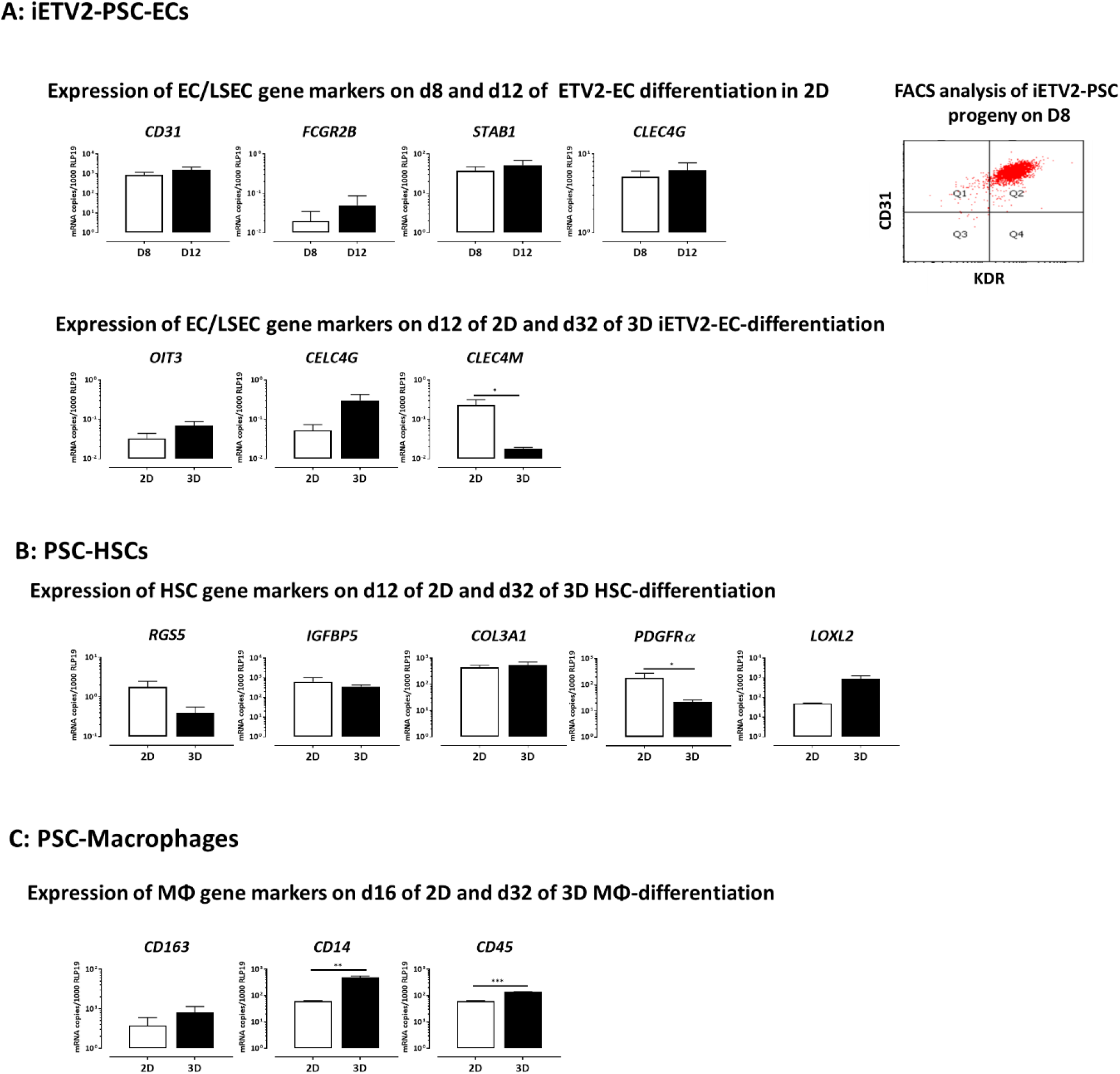
HepMat hydrogel composition also maintains PSC non-parenchymal cells for at least 32 days. (A) PSC-endothelial cells (ECs): iETV2-PSCs were differentiated in 2D to ECs, as described in Fig.3. RT-qPCR for EC/LSEC marker genes (*CD31, FGFR2B, STAB1, CLEC4G*) on d8 and d12 of 2D differentiation (N=3 biological replicates). FACS analysis for CD31 and KDR on day 8 of 2D culture (representative for N=2 biological replicates). RT-qPCR for the LSEC marker genes (*OIT3, CLEC4G* and *CLEC4M*) in ETV2-progeny harvested after 12 days of differentiation in 2D cultures or recovered after an additional 32 days of culture in HepMat hydrogel (N=3 biological replicates). **(B) PSC-hepatic stellate cells (HSCs):** RT-qPCR for HSC marker genes (*RGS5, IGFBP5, PDGFRα, COL3A1* and *LOXL2)* in PSC-HSCs after 12 days of differentiation in 2D cultures or 12 days in 2D culture and additional 32 days of culture in HepMat hydrogel (N=3 biological replicates). **(C) PSC-macrophages (Mϕs):** RT-qPCR for Mϕ marker genes (*CD168, CD14, CD45)* in PSC-Mϕ after 16 days of differentiation in 2D culture or 16 days in 2D culture and 32 days of culture in HepMat hydrogel (N=3 biological replicates). Data shown as mean ± SEM and analyzed by two-tailed student t -test. **p <0.01; ***p <0.001; ****p <0.0001.

**Figure S4.**
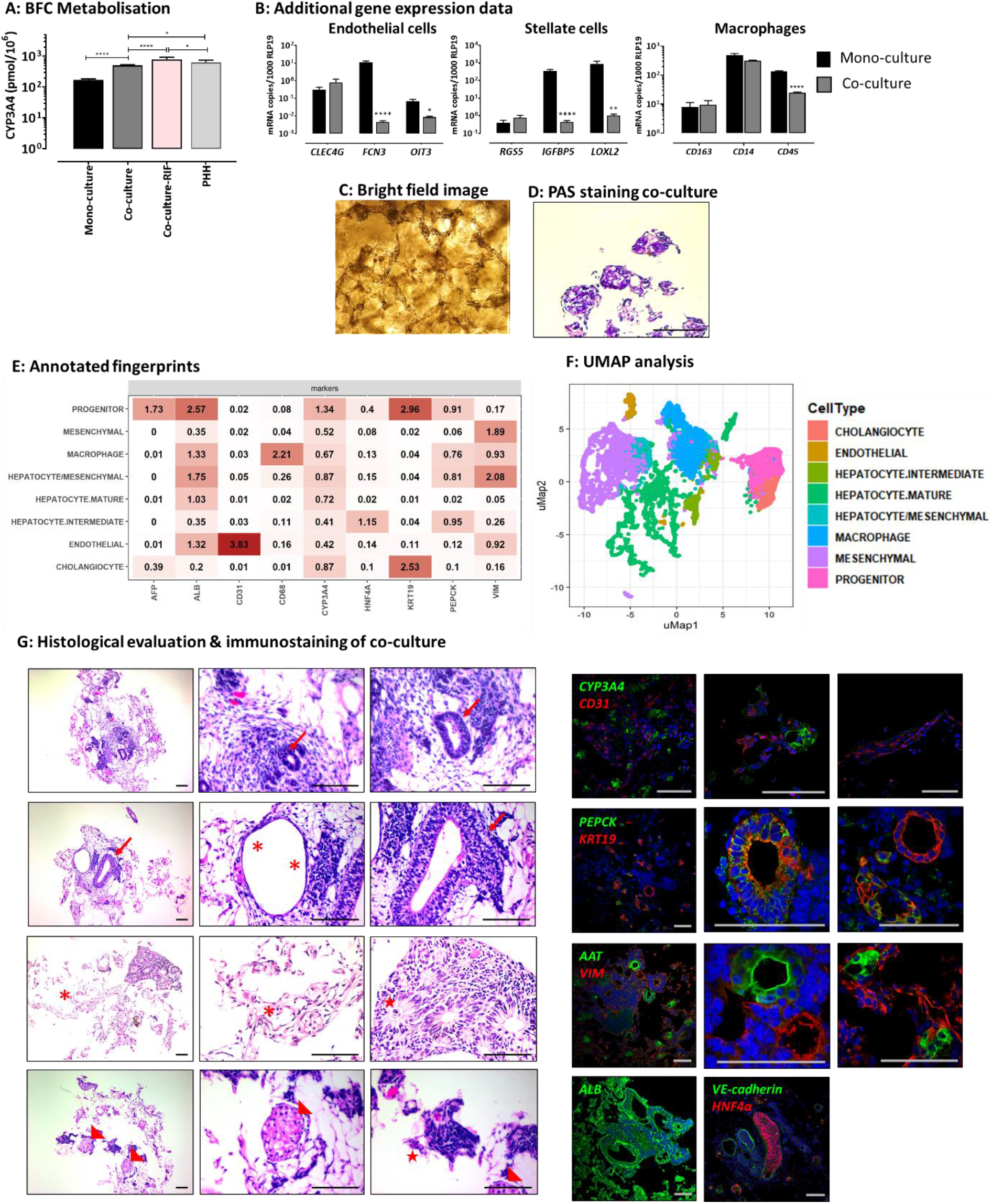
The HepMat hydrogel maintains and supports maturation /quiescence of PSC-hepatocyte-like cells, -endothelial cells, -hepatic stellate cells and -macrophages for at least 32 days. CYP3A4 function defined by BFC metabolization (3D = monoculture of HC3x-HLCs in HepMat hydrogel and liver differentiation medium, N=4; co-culture and co-culture-RIF = HC3x-PSC, ETV2-PSCs, PSC-HSCs and PSC-Mϕ progeny co-culture without and with 25μM of Rifampicin (RIF) for 48h, N=15 and N=6, resp.; PHH; N=3); N = biological replicates)(Data for PHH is the same as in Fig. 2). **(B)** Additional RT-qPCR data for marker genes for the different cell types:*CLEC4G, FCN3* and *OIT3 a*s general EC or LSEC markers, *RGS5, IGFBP5 and LOXL2* as PSC-HSC markers and *MARCO, CD5L, SIGLEC1* and *CD68* PSC-Mϕ/KC markers in cells harvested after 32 days from respective 3D mono-cultures (3D) or 3D co-cultures (co-culture)(N=3 biological replicates (Monoculture data used in this figure is same as in Fig. S2 and Fig.3)).**(C)** Bright field image of co-culture of HC3x-HLC, ETV2-EC, PSC-HSC and PSC-Mϕ progeny after 32 days of culture in HepMat hydrogels. **(D)** Periodic-Acid Schiff (PAS) staining (purple color) of d32 co-culture progeny. **(E-F)** MILAN analysis: expression level per cell group for each of the included markers in multiplex analysis (E); Uniform Manifold Approximation and Projection (UMAP) to perform dimension reduction of all marker used in the MILAN analysis (F). **(G)** Histology and immunostaining of d32 HepMat co-cultures: left panel: H & E staining of paraffin embedded sections; Arrow: cells resembling biliary ducts; Asterix: cells resembling blood vessels; Arrowhead: hepatocyte-like cells. Stars: spindle-shaped stromal cells. Right panel: Immunostaining for Albumin, HNF4α, CYP3A4, PEPCK, KRT19, CD31, VE-cadherin, AAT and Vimentin (representative example of N= 3).

**Figure S5.**
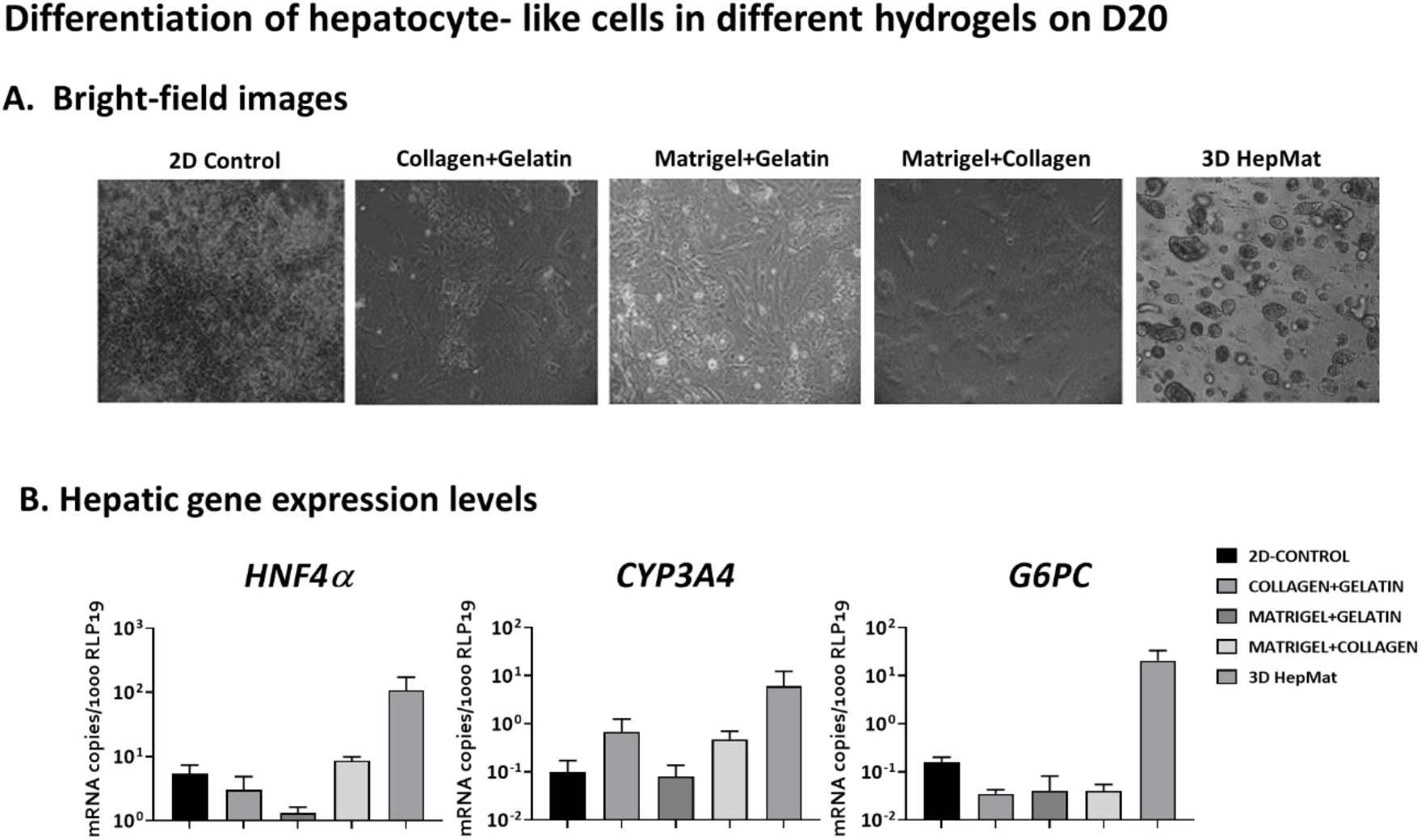
PSC-HLC progeny cultured in different combinations of natural matrices compared with HepMat hydrogel. Day 8 PSC-hepatoblasts were embedded in different natural matrix-based hydrogels, or HepMat hydrogel, and maintained in LDM for an additional 12 days. (A) Representative bright-field images of cultures on D20, showing attachment of PSC progeny cultured in natural matrix hydrogels to the bottom of the wells, which is not seen for cells embedded in the HepMat hydrogel. RT-qPCR for hepatocyte marker genes (*HNF4α, CYP3A4* and *G6PC*) on d20 of differentiation (N=2 biological replicates)

## STAR Methods

### PEG, peptides and hydrogel formation

Vinylsulfone-functionalized four-arm PEG (four-arm PEG-VS 10K) was purchased from Jenkem, USA. The monocysteine ECM/CAM mimicking peptides (Ac-GCGYGXXXG-NH2) and MMP-sensitive crosslinker peptides (Ac-GCRD(E)XXXXX(E)DRCG-NH2) were purchased from Genscript, USA. PEG hydrogels were prepared as described elsewhere with slight modifications (Lutolf and Hubbell, 2003). Four-arm PEG vinyl sulfone was conjugated with monocysteine ECM/CAM peptides (250 µM) via a Michael-type addition reaction in 0.3 M HEPES buffer (pH 7.6) at 37 °C for 30 min to prepare peptide–PEG precursor solution. The peptide–PEG precursor solution was mixed with cells at a concentration of 3×10^7^ cells/ml. Hydrogel formation was triggered by the addition of MMP-sensitive crosslinker peptides mixed in stoichiometrically balanced ratio to generate hydrogel network at 37 °C for 30 min. Dissociation and release of cellular aggregates grown in PEG for downstream cell processing or re-embedding was accomplished by degrading hydrogels with Collagenase Type-1 (Worthington Biochem).

### Screening studies

Three properties of the hydrogels were tuned to create unique 216 microenvironments for testing optimal hepatocyte-like cell (HLC) maturation: (1) combination of six different ECM/CAM peptides listed in Table 1 (Genscript), (2) three different MMP-sensitive crosslinker peptides to tune cell mediated degradability (Genscript), and (3) and three different PEG polymer concentrations to generate different hydrogel stiffnesses. Using the discrete numeric design module in JMP Pro software (SAS, USA), we created 24 unique combinations of six ECM/CAM peptides as shown in Table 2, which in further combination with the different MMP-sensitive crosslinker and different mechanical properties resulted in 216 microenvironments that were tested. The initial screen was done with genetically unmodified PSC-hepatoblasts. Specifically, PSC cells were differentiated until day 8 as described (Boon et al., 2020), at which time they were encapsulated into hydrogels and allowed to differentiate for an additional 8 or 32 days. Assessment of hepatic differentiation was done by RT-qPCR (list of genes can be found in table 3; list of primers can be found in table 5) and -benzyloxy-4-trifluoromethylcoumarin (BFC)-metabolization studies.

**Table 4.** Growth factors/cytokines/additives for co-culture differentiation

| Medium | Cytokines |  |  |
| --- | --- | --- | --- |
|  | Day 0-4 | Day 4-8 | Day 8-32 |
| 50-50% mixture of LDM and StemPro™-34 SFM + Retinol + Palmitic acid | aFGF (20 ng/mL) | HGF (20 ng/mL) | HGF (20 ng/mL) |
|  | bFGF (10 ng/ml) | bFGF (10 ng/ml) | bFGF (10 ng/ml) |
|  | VEGF (50 ng/mL ) | VEGF (50 ng/mL ) | VEGF (50 ng/mL ) |
|  | ECGS (20µl/ml) | ECGS (20µl/ml) | ECGS (20µl/ml) |
|  | MCSF (50 ng/mL) | MCSF (50 ng/mL) | MCSF (5 ng/mL) |
|  | FLT3 (50 ng/mL ) | FLT3 (50 ng/mL ) | 2% Fetal bovine serum |
|  | GM-CSF (25 ng/mL) | GM-CSF (25 ng/mL) |  |
|  | 2% Fetal bovine serum | 2% Fetal bovine serum |  |

**Table 5.**
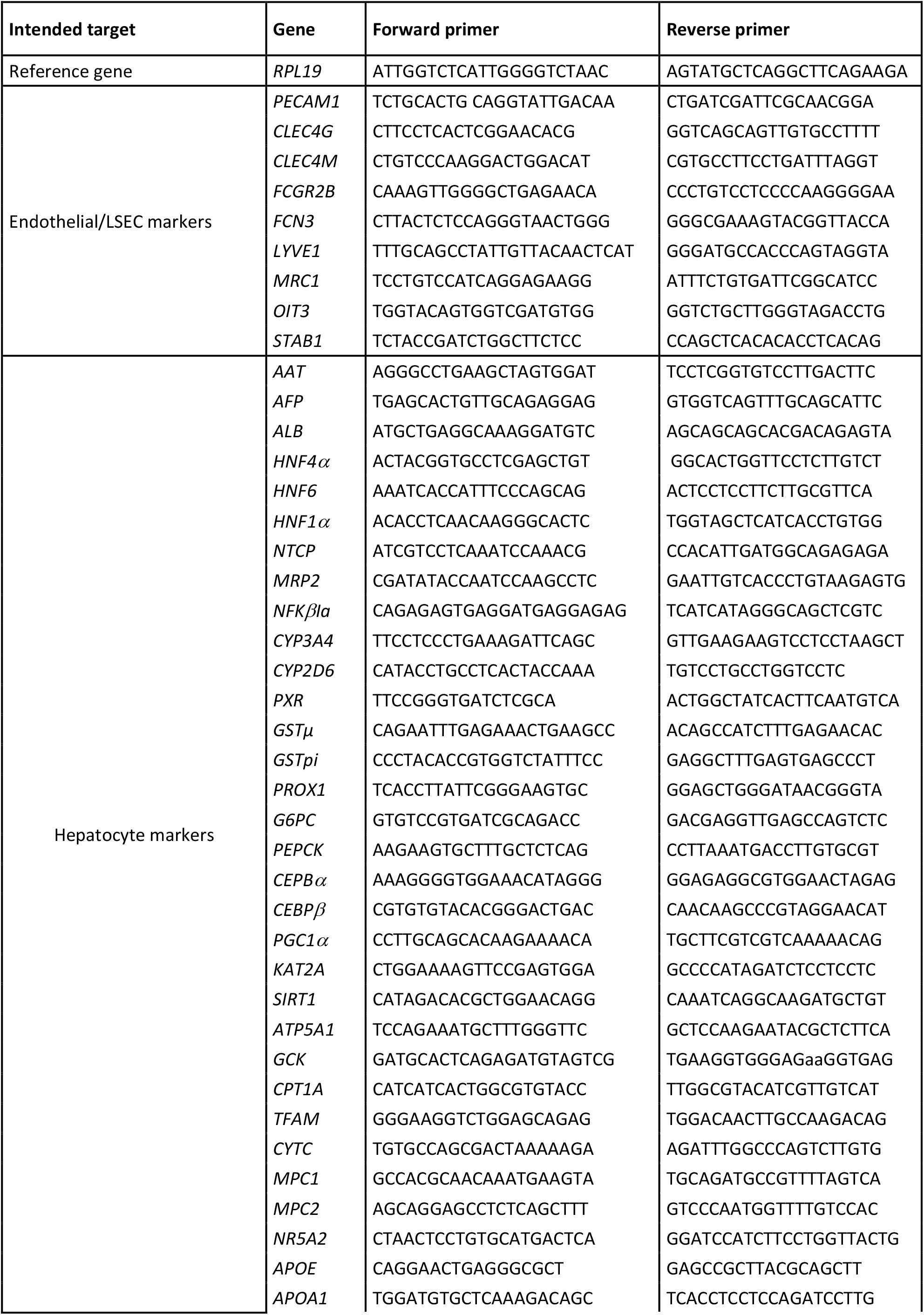
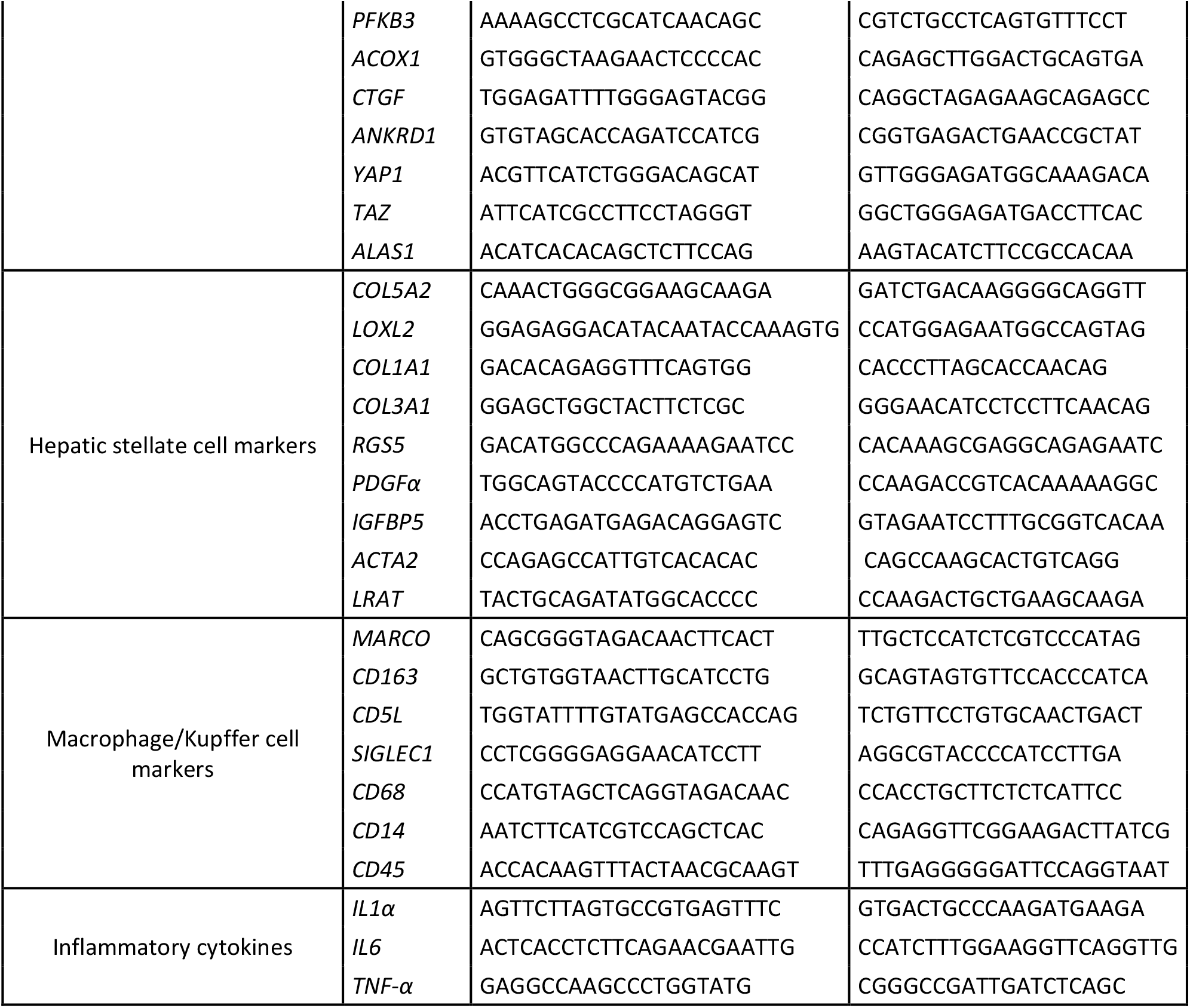
RT-qPCR primer sequences

### Mechanical characterization of PEG hydrogels

The stiffness of PEG hydrogels was determined using a Chiaro Nanoindenter (Optics11, Amsterdam, Netherlands), by applying serial indentations with a spherical glass probe (r, 24.5 µm) attached to a flexible cantilever (k, 0.063 N/m). Loading and unloading velocities of the probe were set to 1.5 and 15 µm/s, respectively, by applying 2 seconds of holding period in between. For each condition, matrix scans (6 × 6 points) from two random locations were obtained from replicate hydrogels. Load vs. displacement curves were extracted individually for each indentation point and Elastic Moduli (E) were calculated by using the Hertzian Contact Model (Poisson’s ratio, 0.5) using the Piuma Dataviewer Software (Optics11).

### Testing the efficiency of peptide conjugation

The efficiency of Michael-type addition reaction performed to generate peptide-PEG precursor solutions was tested by replacing ECM/CAM peptides with a monocysteine 5-FAM peptide (Genscript), at different concentrations (250, 1000 and 2000 μM). After the formation of hydrogels with a DG2 crosslinker peptide, samples were imaged using a LSM 880 Confocal microscope (Zeiss). *Z*-stack of 10 planes from three random locations was scanned for each condition, and average fluorescent intensity was calculated by ImageJ Software.

### hESC differentiation to hepatocyte-like cells

The hESC line H9 (WA09) was purchased from WiCell Research Institute (Madison, 15 WI) and expanded feeder free on matrigel (BD Biosciences) coated plates in Essential 8 or Essential 8 Flex (Thermo Fisher Scientific). All studies using H9 ESCs were approved by the Human Ethics committee at the University Hospital, Gasthuisberg, KU Leuven, Belgium. H9 cells were differentiated towards HLCs as described (Boon et al., 2020). Briefly, H9 cells were made single cell using StemPro™Accutase™ Cell dissociation Reagent (Thermo Fisher Scientific) and plated on Matrigel-coated plates at ±8.75 × 10^4^ cells/cm^2^ in mTeSR medium (Stem Cell Technologies) supplemented with RevitaCell (Thermo Fisher Scientific). When cells reached 70-80% confluence differentiation was started using the previously described cytokine regimens in liver differentiation medium (LDM) and was stopped after 20 or 40 days of differentiation. All cytokines were purchased from Peprotech. Differentiation medium was supplemented with 0.6% dimethylsulfoxide (DMSO) during the first 12 days of the culture and with 2.0% DMSO during the last 8 days of differentiation. The hepatic differentiation from genetically modified PSCs (termed HC3x as in (Boon et al., 2020) was performed in liver differentiation medium (LDM) until D12; after which 3x concentrate of non-essential amino-acids (AAs) was added to the culture until day 14, and from day 14 until the end of the culture, glycine, at a concentration of 20g/L was added combined with the AAs (Boon et al., 2020). When AAs were added, DMSO was omitted from the culture.

### hESC differentiation to ETV2-inducible endothelial cells (iETV2-ECs)

The iETV2 cell line was generated by recombining a Tet-inducible cDNA for *ETV2* in the FRT-flanked cassette in the *AAVS1* locus of PSCs as described (Ordovas et al., 2015). PSCs containing the inducible overexpression cassette for *ETV2* were differentiated towards endothelial cells (ECs) using LDM containing 5 μl/ml doxycycline and 10 ng/ml bFGF starting on day 0 of differentiation. From day 2 onwards, 2.0% fetal bovine serum (FBS) was added to the medium. iETV2-ECs were dissociated with StemPro™Accutase™ Cell dissociation Reagent (Thermo Fisher Scientific) and passaged every 4 days until day 12, when they were encapsulated in hydrogels.

### hESC differentiation to hepatic stellate cell-like cells

Differentiation of PSCs towards hepatic stellate cells (HSCs) was performed as in (Coll *et al*., 2018). H9 cells were grown on Matrigel coated plates until confluency, and then collected as single cells by StemPro™Accutase™ treatment, and plated on matrigel-coated plates at 5 × 10^4^ cells/cm^2^ density in mTeSR medium with RevitaCell Supplement (Thermo Fisher Scientific). Differentiation was started when cells reached 70-80% confluency. At the start of differentiation, mTeSR medium was replaced by LDM containing different cytokine mixes as described by Coll *et al*., 2018. On day 8, cells were harvested with 0.05% Trypsin (Thermo Fisher Scientific) and re-plated with RevitaCell. On day 12, cells were collected by StemPro™Accutase™ treatment and encapsulated in the hydrogels.

### hESC differentiation to macrophages

hESC were differentiated towards macrophages as described in (van Wilgenburg et al., 2013). PSC were resuspended at a final cell concentration of 1×10^5^ cells/mL in mTeSR™-1 spin-EB medium (mTeSR™-1, Stem Cell Technologies), 1 mM Rock-inhibitor (Calbiochem); BMP-4 (50 ng/ml, Peprotech), SCF (20 ng/mL, Peprotech) and VEGF (50 ng/mL, Peprotech). 100 µL was added per well of 96-well ultra-low adherence plates (Greiner Bio-one). Plates were centrifuged at 300 RCF for 5 min, and incubated for 4 days at 37°C and 5% CO_2_. EBs were fed every day by gently aspirating 50 µL medium and gently adding 50 µL of fresh EB medium. On day 4, approximately 20 EBs were transferred into one well of a six-well tissue culture plate in 4 mL medium (X-VIVOTM15 (Lonza)) supplemented with glutamax (2 mM, Invitrogen), SCF (50 ng/ml, Peprotech), M-CSF (50 ng/ml, Peprotech), IL3 (50 ng/ml), FLT3 (50 ng/ml, Peprotech) and TPO (5 ng/ml, Peprotech) 100 U/mL penicillin,100 µg/mL streptomycin (Invitrogen) and β-mercaptoethanol (0.055 mM, Invitrogen) until day 11 with a media change on day 8. From day 11 onwards, X-VIVOTM15 (Lonza) supplemented with glutamax (2 mM,) β-mercaptoethanol (0.055 mM, Invitrogen), FLT3 (50 ng/ml), M-CSF (50 ng/ml) and GM-CSF (25 ng/ml, Peprotech) till the end of the differentiation with a media change every week. From day 16 onwards, cells were collected for encapsulation in hydrogels either for mono-culture or co-culture studies.

### Co-cultures

The different PSC-liver cell progeny were encapsulated in PEG hydrogels at different time points. Hepatoblasts were harvested on D8; iETV2-ECs on D12; HSCs on day 12 and Mϕs from day 16 onwards, and encapsulated in HepMat hydrogel at a ratio of 2:1:1:0.5, respectively. The media used for these co-culture experiments was a combination of LDM and StemPro™-34 SFM medium (1:1) combined with all growth factors/cytokines/additives used for each cell type. The co-culture medium consisted therefore of: 50:50% LDM and Stem Pro 34, as well as Retinol + Palmitic acid, and a combination of cytokines/growth factors as described in Table 4). Encapsulated cells were maintained and differentiated in hydrogels for an additional 32 days.

### RNA extraction and reverse-transcription quantitative PCR (RT-qPCR)

RNA extraction was performed using TRIzol reagent (Invitrogen) following manufacturer’s instructions. At least 1µg of RNA was transcribed to cDNA using the Superscript III First-Strand synthesis (Invitrogen). Gene expression analysis was performed using the Platinum SYBR green qPCR supermix-UDG kit (Invitrogen) in a ViiA 7 Real-Time PCR instrument (Thermo Fisher Scientific). The sequences of all used RT-qPCR primers are listed in Table 5. The ribosomal protein L19 transcript (*RPL19*) was used as a housekeeping gene for normalization.

### Histology

Hydrogels were fixed with 4% (w/v) paraformaldehyde (PFA, Sigma-Aldrich) overnight at 4°C, washed 3 times with PBS and submerged in PBS-sodium azide (0.01% v/v) solution at 4°C until embedded in paraffin. Hydrogel sections (5 µm) were prepared using a microtome (Microm HM 360, Marshall Scientific.)

For Hematoxylin and Eosin (H&E) and Periodic-acid Schiff (PAS) staining, sections were treated with xylene solution to remove the paraffin, and gradually rehydrated in ethanol (100 to 70%, v/v). H&E staining was performed by submerging rehydrated hydrogel sections in Harris Hematoxylin solution, acid alcohol, bluing reagent and Eosin-Y solution by order. Stained samples were dehydrated with ascending alcohol series, washed in xylene solution, and mounted with DPX mountant (Sigma-Aldrich). PAS staining were performed according to the manufacturer’s instructions.

### Immunofluorescence analysis

Following deparaffinization in xylene and rehydration in descending alcohol series, heat-mediated antigen retrieval was performed by incubating hydrogel sections in Dako antigen retrieval solution (Dako) for 20 min at 98 °C. This step was followed by cell permeabilization with 0.01% (v/v) Triton-X (Sigma-Aldrich) solution in PBS, for 20 minutes. Samples were then incubated with 5% (v/v) Goat or Donkey Serum (Dako) for 30 min. Primary antibodies (Table 6) diluted in Dako antibody diluent solution, were incubated overnight at 4°C, followed by washing steps and incubation with Alexa-coupled secondary antibody (1:500) and Hoechst 33412 (1:500) solution for 1 hour at room temperature. Finally, samples were washed in PBS, and mounted with Vectashield antifade mounting medium (Vector Laboratories). Stained sections were imaged using laser scanning confocal microscope (LSM 880, Zeiss, Germany), and image processing were performed on ZEN Blue software (Zeiss, Germany).

**Table 6.** List of antibodies

| ID | Antibody/Dye | Catalogue number | Company | Dilution | Application |
| --- | --- | --- | --- | --- | --- |
| 1 | Anti-CD144 (VE-cadherin) | 13-1449-82 | Thermo Fisher Scientific | 1/50 | Flow cytometry |
| 2 | Anti-VE Cadherin | ab33168 | Abcam | 1/150 | Immunostaining |
| 3 | Anti-HNF4α | ab41898 | Abcam | 1/200 | Immunostaining |
| 4 | Anti-SLC10A1 | HPA042727 | Sigma-Aldrich | 1/500 | Immunostaining |
| 5 | Anti- α1-Antitrypsin | A0012 | DAKO | 1/200 | Immunostaining |
| 6 | Anti-YP3A4 | PAP 011 | Tebu-Bio (Cypex limited (Dundee, UK)) | 1/250 | Immunostaining |
| 7 | Anti-PEPCK (H-300) | sc-271204 | Santa Cruz | 1/1000 | Immunostaining |
| 8 | Anti-CK19 | sc-33120 | Santa Cruz | 1/500 | Immunostaining |
| 9 | Anti-Vimentin | M0725 | Dako | 1/500 | Immunostaining |
| 10 | Anti-PECAM1 | M0823 | Dako | 1/20 | Flow cytometry |
| 11 | Anti-PECAM1 | ab33168 | Abcam | 1/200 | Immunostaining |
| 12 | Anti-CD68 | MA5-12407 | Thermo Fisher Scientific | 1 μg/ml | Immunostaining |
| 13 | Anti-MRP2 | ab3373 | Abcam | 1/500 | Immunostaining |
| 14 | Anti-Occludin | ab31721 | Abcam | 1/500 | Immunostaining |
| 15 | Anti-AFP (C-19) | MAB1368 | R&D Systems | 1/500 | Immunostaining |
| 16 | Anti-CD45 | ab10558 | Abcam | 1/150 | Flow cytometry |
| 17 | Donkey anti-Mouse IgG (H+L)<br>Alexa Fluor 488 | A-11029 | Molecular probes | 1/500 | Immunostaining/<br>Flow cytometry |
| 18 | Donkey anti-Rabbit IgG (H+L)<br>Alexa Fluor 555 | A21429 | Molecular probes | 1/500 | Immunostaining/<br>Flow cytometry |
| 19 | Donkey anti-Rabbit IgG (H+L)<br>Alexa Fluor 647 | A-31573 | Thermo Fisher Scientific | Use as control | Immunostaining/<br>Flow cytometry |
| 20 | Rabbit IgG | 550875 | BD Pharmingen | Used as control | Immunostaining |
| 21 | Mouse BALB/c IgG1 | 550878 | BD Pharmingen | Used as control | Immunostaining |
| 22 | eBioscience™ Streptavidin<br>eFluor™ 450 Conjugate | 48-4317-82 | Thermo Fisher Scientific |  | Flow cytometry |
| 23 | Alexa Fluor™ 488 Phalloidin | A12379 | Thermo Fisher Scientific | 2 units/ml | Immunostaining |
| 24 | Prolong Gold antifade reagent<br>with DAPI | P-36931 | Thermo Fisher Scientific |  | Immunostaining |
| 25 | Hoechst 33342 | H3570 | Thermo Fisher Scientific | 1/500 | Immunostaining |
| 26 | Calcein-AM | C1430 | Thermo Fisher Scientific | 2 μM | Staining |
| 27 | Ethidium homodimer-1 | C1430 | Thermo Fisher Scientific | 2 μM | Staining |
| 28 | Propidium iodide | 81845 | Sigma-Aldrich | 100ng/ml | Flow cytometry |

### Multiplex-stripping immunofluorescence and Image pre-processing

The multiplex staining was performed according to the MILAN protocol previously described by (Bolognesi et al., 2017; Bosisio et al., 2020). MILAN entails multiple rounds of indirect immune-fluorescent staining, imaging and antibody removal via a detergent and a reducing agent. Unconjugated primary antibodies (see Table 5) were used. Following the MILAN protocol, custom pipelines implemented in R software (versions 3.5 and 3.6) were used to pre-process the image. Image registration was performed by applying a homomorphic transformation over a set of matched descriptors using a Harris detector in the DAPI channel for different rounds of staining. Registration was followed by autofluorescence (AF) subtraction, previously acquired. AF subtraction was performed by subtracting the scaled AF channel from the measured signal (MS). Cell segmentation was performed using the Intelesis plug-in (from Zeiss). Quality Control (QC) over the Mean Fluorescence Intensity (MϕI) values was performed by evaluating the signal-to-noise-ratio (SNR) of every signal. Three independent clustering methods were used to group cells with similar expression profiles. Each cluster was characterized by the average expression of its cells for each marker. For each clustering method, clusters were manually annotated to known cell types by expert pathologists (FMB, MVH, double blinded analysis). Only cells agreeing in 2 or more clustering methods were considered robust and mapped to known cell phenotypes. The rest of the cells were labelled as BLANK.

### Flow cytometry

Cells present in HepMat cultures were isolated by collagenase treatment of 6 hydrogels. Isolated clusters were washed with 1 x PBS and dissociated to single cells by treatment with TrypLE™ Express Enzyme (1X), phenol red (Thermo Fisher Scientific) for 20 min. After a PBS wash, single cells were stained with the primary antibodies for 45 min along with respective isotype controls at the concentration shown in Table 6. Dead cells were excluded by propidium iodide staining. All samples were analyzed using a FACS-Canto (BD Biosciences).

### Functional assessments

#### Hepatic progeny

To assess glycogen storage, we stained sections using periodic acid-Schiff (PAS, Sigma-Aldrich). CYP3A4 dependent metabolization over 4h was determined using the fluorimetric probe BFC. Albumin secretion rate was quantified using the human albumin ELISA quantitation kit (Bethyl Laboratory).

#### TGFβ exposure of HepMat-PSC-HSC cultures

Day 32 HepMat-PSC-HSC mono-cultures hydrogels were exposed to 25 ng/ml of TGFβ (Peprotech) on day 32. Expression of inflammatory and fibrogenic genes was determined by RT-1PCR. Supernatants were collected for pro-collagen measurement by ELISA.

#### LPS exposure of HepMat-PSC-Mϕs cultures

Day 32 HepMat-PSC-Mϕs mono-cultures were exposed to 100 ng/ml of LPS (Sigma-Aldrich). Expression of inflammatory genes was determined by RT-qPCR. Supernatants were collected for inflammatory cytokine measurement by ELISA.

#### TGFβ exposure of HepMat-co-cultures and mono-cultures

Day 32 HepMat co-cultures and dayy 32 HepMat HC3x HLC, HSC and Mϕ mono-cultures were incubated with 25 ng/ml TGFβ for 7 days (addition on day 0 and day 3). Expression of inflammatory and fibrogenic genes was determined by RT-qPCR. Supernatants were collected for pro-collagen measurement by ELISA.

#### Oleic acid (OA) exposure of HepMat-co-cultures and mono-cultures

Day 32 HepMat co-cultures and Day 32 HepMat HC3x HLC, HSC and Mϕ mono-cultures were incubated with 800μM of OA (Sigma-Aldrich) for 7 days (addition on day 0 and day 3). Day 32 HepMat co-cultures were also incubated with a combination 800μM of OA and 1μM Obeticholic acid (OCA) (MedChemExpress) or 800μM of OA and 30µM Elafibranor (ELN) (MedChemExpress) for 3 days (addition on day 0 only). Expression of inflammatory and fibrogenic genes was determined by RT-1PCR. Supernatants were collected for pro-collagen measurement by ELISA on day 3 and day 7. In addition, co-culture were assayed for presence of steatosis by BODIPY® staining on day 3 and day 7 as follows: Gels were transferred to Glass Bottom plates (Cellvis) for live-cell imaging, following incubation with BODIPY® 493/503 for lipids (Thermo Fisher Scientific) and Hoechst 33342 (Sigma-Aldrich). Sample was visualized and scanned on a laser scanning confocal microscope (LSM 880, Zeiss).

### TNF-α, IL1α, IL6 and Pro-collagen 1 ELISA

The supernatant was assayed for human TNF-α, IL1α, and IL6 by ELISA (Biolegend), according to the manufacturer’s instructions. Presence of pro-collagen type I was detected by Pro-collagen Type I C-peptide ELISA kit (Takara Bio Inc). Secreted level of TNF-α, IL1α, IL6 and pro-collagen 1 were normalize the for the cell number.

### Quantification and Statistical Analysis

Results are expressed as the arithmetic mean ± standard error of the mean (SEM). All experimental results are from a minimum of 3 biological replicate experiments unless otherwise stated. Statistical comparisons between groups were done using Student’s t-test, one way Anova (Tukey’s multiple comparison) test when appropriate. A *p-value* of < 0.05 was considered significant. Analyses were carried out using either JMP pro software (SAS Institute) or GraphPad Prism 8.0 (GraphPad prism Software Inc.).

